# Coordinated post-transcriptional control of oncogene-induced senescence by UNR/CSDE1

**DOI:** 10.1101/2020.12.08.415794

**Authors:** Rosario Avolio, Marta Inglés-Ferrandiz, Annagiulia Ciocia, Sarah Bonnin, Anna Ribó, Fátima Gebauer

**Affiliations:** Gene Regulation, Stem Cells and Cancer Programme, The Barcelona Institute of Science and Technology, 08003 Barcelona, Spain; Bioinformatics Unit, Centre for Genomic Regulation (CRG), The Barcelona Institute of Science and Technology, 08003 Barcelona, Spain; Universitat Pompeu Fabra (UPF), 08003 Barcelona, Spain; Department of Molecular Medicine and Medical Biotechnology, University of Napoli “Federico II”, 80131 Napoli, Italy; Biocruces Bizkaia Health Research Institute, Research Unit of Basque Center for Blood Transfusion and Human Tissues, 01004 Bizkaia, Spain

**Author notes:** Equal contribution (Rosario Avolio, Marta Inglés-Ferrandiz).

## Abstract

Oncogene-induced senescence (OIS) is a form of stable cell cycle arrest elicited in cells as a response to oncogenic stimulation. OIS must be bypassed for transformation, but the mechanisms of OIS establishment and bypass remain poorly understood, especially at the post-transcriptional level. Here we show that the RNA binding protein UNR/CSDE1, previously involved in melanoma metastasis, unexpectedly enables OIS in primary mouse keratinocytes that have been challenged by over-expression of oncogenic H-Ras. Depletion of CSDE1 leads to senescence bypass, cell immortalization and tumor formation in vivo, indicating that CSDE1 behaves as a tumor suppressor. Using iCLIP-Seq, RNA-Seq and polysome profiling we have uncovered two independent molecular branches by which CSDE1 contributes to OIS. On one hand, CSDE1 enhances the senescence-associated secretory phenotype (SASP) by promoting the stability of SASP factor mRNAs. On the other hand, CSDE1 represses the synthesis of the pro-oncogenic RNA binding protein YBX1. Importantly, depletion of YBX1 from immortal keratinocytes rescues senescence and uncouples proliferation arrest from the SASP, revealing multilayered post-transcriptional mechanisms exerted by CSDE1 to control senescence. Our data uncover a novel function of CSDE1, and highlight the relevance of post-transcriptional control in the regulation of senescence.

## INTRODUCTION

Oncogenic transformation is a multistep process usually initiated by aberrant activation of driver oncogenes, such as the members of the Rat sarcoma (Ras) GTPase family (including K-Ras, N-Ras and H-Ras) (Hanahan and Weinberg, 2011). In response to oncogenic activation, cells can adopt counter-acting mechanisms as a first line of defense against unrestricted proliferation. One such mechanism is known as oncogene-induced senescence (OIS) (Zhu et al., 2020). OIS was described for the first time in 1997, when Serrano et al. showed that expression of oncogenic H-Ras^V12^ in primary fibroblasts induces a stable cell cycle arrest accompanied by morphological changes similar to those occurring during cellular replicative senescence (Serrano et al., 1997). Among other features, senescence includes the secretion of a cocktail of cytokines, chemokines, growth factors and proteases collectively known as the senescence-associated secretory phenotype (SASP) (Gorgoulis et al., 2019; Herranz and Gil, 2018). While the exact composition of the SASP varies with the senescence trigger and the cell type, it exerts autocrine and paracrine effects, reinforcing senescence in the cell of origin and the surrounding microenvironment (Basisty et al., 2020).

The relationship of senescence and the SASP with tumorigenesis is complex. SASP factors can potentiate immunosurveillance, guiding the immune system to build anti-cancer responses, but they can also enhance immunosuppression and promote the escape of metastatic cells (Acosta et al., 2013; Rao and Jackson, 2016; Ruscetti et al., 2020). Similarly, while early senescence is considered a tumor suppressive mechanism that must be bypassed for oncogenic transformation, prolongued senescence can promote the persistence of therapy-resistant cells and tumor relapse (Wang et al., 2020). Thus, further research is necessary to understand the mechanisms of senescence and their relationships with the oncogenic process, including the identification of new players relevant for the establishment or maintenance of OIS. In this context, post-transcriptional regulation of OIS represents a relatively unexplored field. RNA binding proteins (RBPs) regulate every post-transcriptional step of gene expression, but only a limited number of RBPs have been related to senescence to date (Fregoso et al., 2013; Groisman et al., 2006; Kwon et al., 2018; Majumder et al., 2016; Radine et al., 2020; Sanduja et al., 2009; Wang et al., 2005; Wang et al., 2001). Here we uncover a novel RBP in senescence and unravel its molecular mechanisms of action.

Cold Shock Domain Containing E1 (CSDE1), also termed Upstream of N-Ras (UNR), is a conserved RBP with a high RNA binding capacity endowed by multiple cold shock domains (Anderson and Catnaigh, 2015; Guo et al., 2020; Hollmann et al., 2020; Mihailovich et al., 2010). CSDE1 displays context-specific functions in apoptosis and differentiation (Dormoy-Raclet et al., 2007; Elatmani et al., 2011; Horos et al., 2012; Ju Lee et al., 2017). We have previously shown that CSDE1 behaves as a pro-metastatic factor in melanoma, where it regulates the levels and translation of mRNAs encoding important tumor suppressors (e.g. PTEN) and EMT markers (e.g. VIM, RAC1) (Wurth et al., 2016). While the oncogenic potential of CSDE1 has been observed in multiple other tumors (Fang et al., 2005; Liu et al., 2020; Martinez-Useros et al., 2019; Tian et al., 2018), loss of function CSDE1 mutations have been detected in neuroendocrine tumors, suggesting tumor suppressive roles (Fishbein et al., 2017). Intrigued by context-specific functions of CSDE1 in cancer, we noticed that this RBP could bind SASP factor mRNAs in melanoma, although such binding did not have functional consequences in this system (Wurth et al., 2016). We, thus, wondered whether CSDE1 could have a role in the context of senescence representing earlier steps of transformation. Here we show that CSDE1 is necessary for H-Ras-induced senescence in primary mouse keratinocytes. Depletion of CSDE1 leads to senescence bypass, cell immortalization and transformation, indicating tumor suppressive roles. Unbiased multi-omics studies uncovered two independent mechanisms coordinated by CSDE1 to promote senescence, stabilization of SASP factor mRNAs and inhibition of YBX1 mRNA translation. Notably, transformed keratinocytes reverted back to a senescent state upon depletion of YBX1, highlighting the relevance of CSDE1-mediated translational control.

## RESULTS

### CSDE1 promotes oncogene-induced senescence

We first assessed whether CSDE1 contributes to the senescence program using a well-established OIS system in primary mouse keratinocytes (PMK) (Ritschka et al., 2017). Briefly, PMK were freshly isolated from the skin of newborn mice and challenged by overexpression of the oncogene *H-Ras^V12^* (referred to as Ras hereafter) using retroviral infection in order to induce senescence. As controls, we infected PMK with an empty retroviral vector (EV) or used non-infected (NI) PMK, and monitored the progression of senescence at various times after induction (Figure 1A).

**Figure 1.**
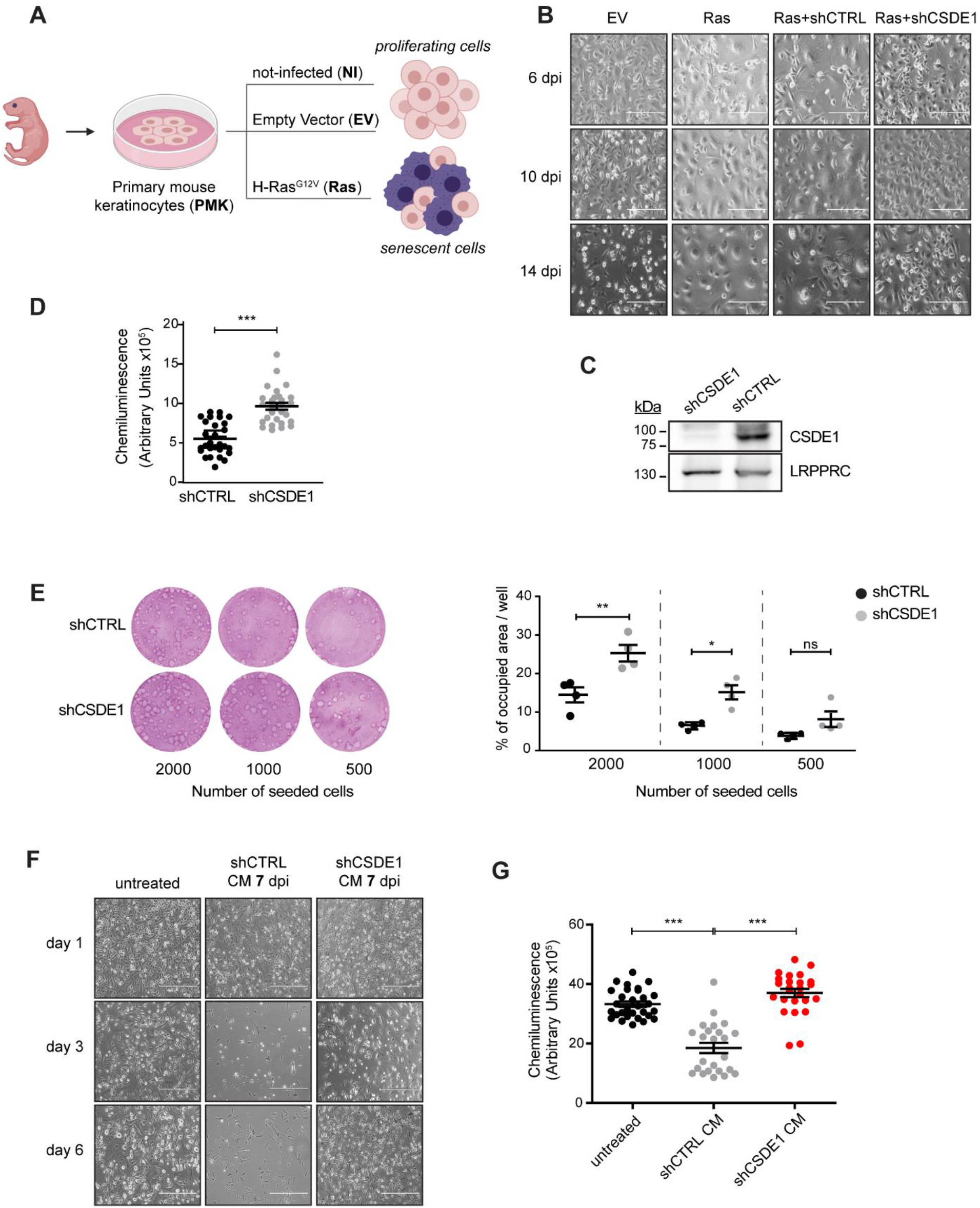
CSDE1 promotes OIS and reinforces the SASP. **(A)** Schematic representation of the OIS experimental model. **(B)** CSDE1 depletion prevents the appearance of senescent cells. Bright field microscopy images of primary mouse keratinocytes (PMK) at various times after infection. dpi, days post-infection; scale bar: 200 μm. **(C)** Efficiency of CSDE1 depletion assessed by Western blot. A representative example of Ras cells at 12 dpi is shown. LRPPRC was used as a loading control. **(D)** CSDE1 depletion leads to increased proliferation, as measured by BrdU incorporation. The experiment was performed at 9 dpi in three biological replicates (n= 3, each with 11 technical replicates). **(E)** CSDE1 depletion results in increased colony-formation capacity. Increasing numbers of shCTRL and shCSDE1 cells were seeded on a layer of mitotically inactivated feeder cells. Quantification is shown on the right (n = 2, each with 2 technical replicates). **(F)** CSDE1 depletion impairs the SASP. Fresh PMK were incubated for the indicated days with media from shCTRL or shCSDE1 cells taken at 7 dpi, using media from non-infected cells (untreated) as control, and visualized by microscopy. CM, conditioned media; scale bar = 400 μm. **(G)** The results in (F) were confirmed by BrdU incorporation assays using a similar number of viable cells after 5 days of CM treatment (n=3, each with 10 technical replicates). Significance in all assays was assessed by unpaired Student’s t-test (*p-value <0.05, **p-value < 0.01, ***p-value < 0.001).

OIS is accompanied by a complete remodeling of gene expression that leads to profound changes in cell morphology and an irreversible cell cycle arrest (Serrano et al., 1997). Because cells grow but do not divide, cells acquire a typical enlarged appearance easily distinguishable under the bright-field microscope (Figure 1B, compare EV with Ras panels). Senescent cells are already evident as early as 6 days post-infection (dpi). To test for a potential effect of CSDE1 in this process, we co-infected Ras-expressing cells with either a CSDE1-targeting shRNA (shCSDE1) or a non-targeting control (shCTRL). Notably, while shCTRL cells were indistinguishable from cells infected with Ras alone, shCSDE1 cells appeared to evade senescence, as judged by a reduced number of enlarged cells and the prevalence of small, proliferating cells (Figure 1B, see efficiency of depletion in Figure 1C). To confirm that shCSDE1 cells indeed bypass senescence, we used a variety of alternative assays. First, we quantified proliferation using BrdU incorporation assays, confirming a higher proliferation rate of shCSDE1 cells compared to shCTRL (Fig 1D).

Second, we performed colony-forming assays, which showed that both the number and size of colonies increased upon CSDE1 depletion (Figure 1E). Finally, as it has been shown previously that chronic exposure to the SASP can induce paracrine senescence (Ritschka et al 2017, Acosta et al 2013), we performed conditioned media (CM) experiments to evaluate whether CSDE1 depletion influenced the SASP. To this end, we collected media from shCTRL and shCSDE1 cells at an early point of senescence (7 dpi) and challenged freshly isolated PMK with these media, carrying medium from untreated PMK as negative control. Significant cell death was already detected at day 3 after incubation with shCTRL-CM media, while cells treated with shCSDE1-CM remained unaffected (Figure 1F). Further, BrdU incorporation assays showed that shCTRL-CM treated cells dramatically reduced proliferation, as expected from an active SASP in the medium, while shCSDE1-CM treated cells proliferated similarly to control cells (Figure 1G).

Taken together, these data indicate that CSDE1 is required for OIS induction and SASP production.

### CSDE1 is a tumor suppressor in primary keratinocytes

Primary cells that bypass OIS may be cultured for longer periods of time, eventually acquiring an immortalized phenotype (Maurelli et al., 2016). Accordingly, depletion of CSDE1 allowed Ras-infected PMK to be extendedly cultured. We have been able to generate three independent shCSDE1 cell lines that have been cultured so far indefinitely, and which we term immortalized PMK (iPMK) (Figure 2A). Western-blot analysis performed at passage 20 confirmed that the lack of CSDE1 is maintained in these cells (Figure 2B). As cell immortalization is a prerequisite for malignant transformation, we next evaluated the tumorigenic potential of the iPMK cells using xenograft experiments. iPMK cells were implanted subcutaneously into immunodeficient mice and tumor growth was monitored for 4-5 weeks. Strikingly, all iPMK cell lines led to tumor formation in 15-20 days, albeit at different growth rates, as shown by measurement of tumor weight at the experimental endpoint (Figure 2C). These results indicate that iPMK cells are transformed, and that CSDE1 behaves as a tumor suppressor in PMK.

**Figure 2.**
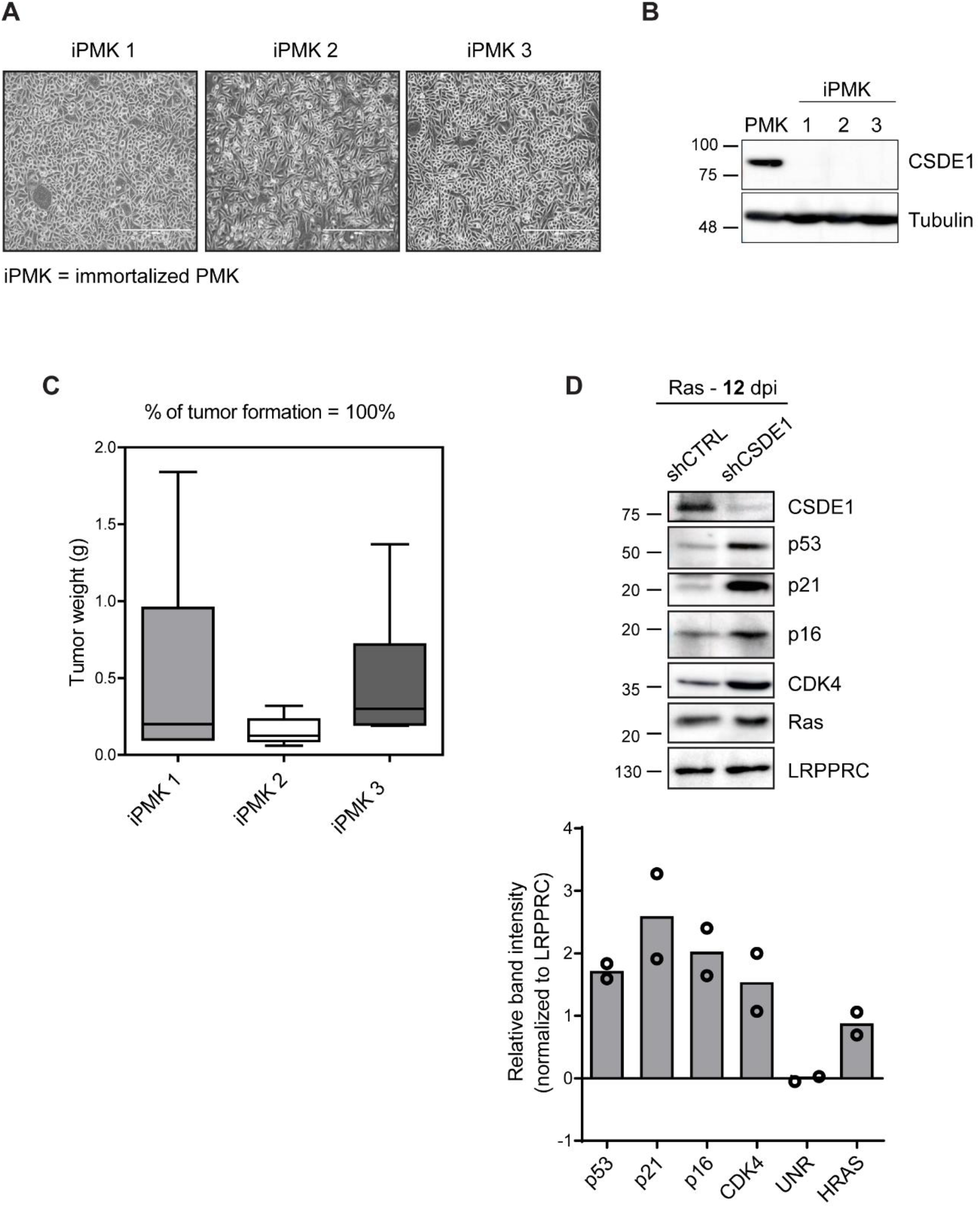
CSDE1 behaves as a tumor suppressor in PMK. **(A)** Bright field microscopy images of three immortalized shCSDE1 cell lines, showing the typical appearance of proliferating cells (iPMK). Scale bar = 400 μm. **(B)** Depletion of CSDE1 is maintained in iPMK. Normal PMK is shown as control. **(C)** iPMK cells are transformed. Cells were injected subcutaneously into nude mice, and tumors were resected and weighted (3 mice were injected in 2 flanks per cell line). **(D)** Expression of senescence markers following CSDE1 depletion, assessed by Western blot. Quantification from two independent experiments is shown at the bottom.

The most parsimonious explanation of the effect of CSDE1 in OIS is that CSDE1 directly promotes the expression of cell cycle inhibitors, such as p53 and its downstream targets p21 and p16, which are typically up-regulated during OIS to mediate cell cycle arrest (Ruiz et al., 2008). To test this hypothesis, we tested the levels of these senescence markers following CSDE1 depletion. Western-blot analysis indicated that none of these markers was downregulated in shCSDE1 cells but, rather surprisingly, they were upregulated compared to control cells (Figure 2D). shCSDE1 cells also show high levels of the proliferation marker CDK4, in agreement with their higher proliferation rate, while the levels of Ras were unaffected (Figure 2D). These results show that CSDE1 depletion leads to senescence bypass despite increased levels of cell cycle inhibitors, resulting in tumorigenesis.

In light of these results, we decided to undertake an unbiased approach to unravel the mechanisms responsible for CSDE1-mediated OIS.

### CSDE1 binds SASP and senescence-related transcripts

To identify CSDE1-bound targets during OIS, we performed iCLIP experiments in proliferating (NI and EV) and senescent (Ras) PMK at an early time of senescence induction (7 dpi) (Figure 3A). Immunoprecipitation of CSDE1 was specific, as the signal was dramatically reduced upon CSDE1 depletion and no signal was detected in the IgG control (Figure S1A). As expected, CSDE1-RNA complexes were detected only upon UV-crosslinking. We performed three independent biological replicates per condition and used non-UV irradiated cells as control (Figure S1B).

**Figure 3.**
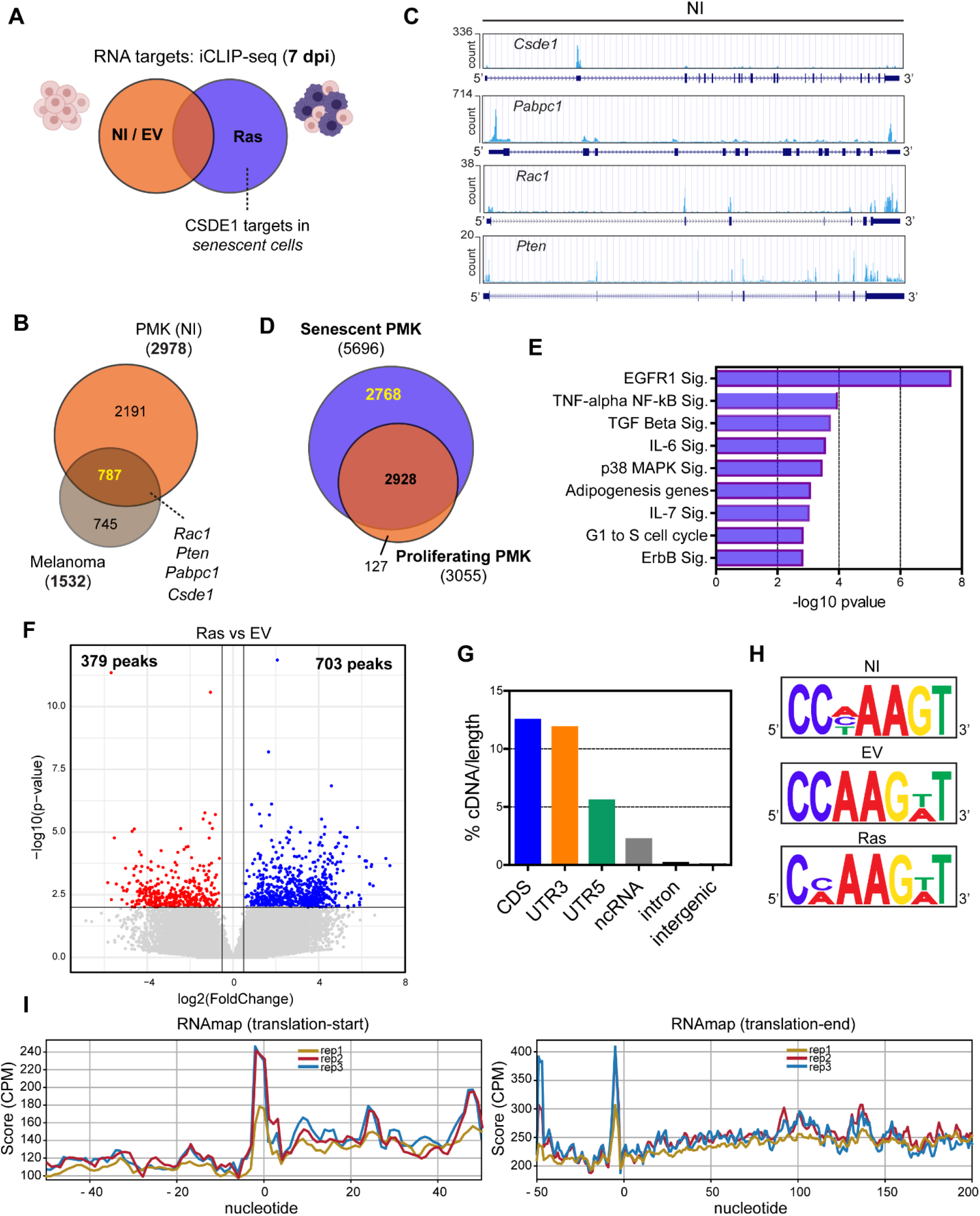
Identification of CSDE1 targets by iCLIP-seq. **(A)** Experimental strategy to identify CSDE1 targets. iCLIP was performed in three independent biological replicates for each condition. **(B)** Overlap between CSDE1 targets in PMK and melanoma cells. **(C)** CSDE1 iCLIP profiles for select genes. **(D)** Overlap of targets bound by CSDE1 in proliferating (pooled NI and EV data) and senescent conditions. **(E)** GO analysis of 2768 targets that are bound de novo in senescent conditions. **(F)** Differential binding of CSDE1 to common sites (peaks) in Ras-vs EV-infected PMK. **(G)** CSDE1 iCLIP density in different transcript regions. Data for non-infected PMK are shown. Results were similar for Ras-and EV-infected cells. **(H)** CSDE1 binding motif in non-infected (NI), EV-and Ras-infected PMK. **(I)** Meta-analysis of CSDE1 binding around the start and stop codons (positioned at 0) in NI cells. Results were similar for Ras-and EV-infected cells.

PCA analysis showed that all replicates belonging to the same condition cluster together and set apart from negative controls (Figure S1C). We considered targets with significant peaks (FDR < 0.05) in the three replicates, and removed those scoring positive in the non-crosslinked controls, which anyway showed a very poor overlap with the crosslinked samples (Figure S1D). The vast majority of the identified targets were protein-coding genes (Table S1). Independent RNA immunoprecipitation (RIP) experiments yielded a validation rate of nearly 100% (Figure S1E).

We have previously identified CSDE1 targets in melanoma cells (Wurth et al., 2016). We thus compared the CSDE1 targets identified in PMK (NI) with those identified in melanoma SK-Mel-147 cells. Around 50% of the targets bound to CSDE1 in both cell types, suggesting conserved functions (Figure 3B). Among them, we find *Csde1* itself, which is known to bind to an IRES located in its 5’-UTR to repress its own translation (Schepens et al., 2007); *Pabpc1*, an important regulatory target of CSDE1(Patel et al., 2005); and curiously also targets functionally relevant for the pro-oncogenic roles of CSDE1 such as *Rac1* and *Pten* (Wurth et al., 2016) (Figures 3B-C). Comparison of proliferating and senescent cells revealed a set of 2768 transcripts binding *de novo* to CSDE1 upon senescence (Figure 3D). Gene ontology (GO) analysis of this subset showed enrichment for pathways related to senescence and SASP signatures, such as EGFR1, TNF-α, TGF-β, IL-6 and MAPK signaling (Figure 3E). Indeed, a prominent group of transcripts bound by CSDE1 consisted of mRNAs encoding SASP factors. We built a list of 522 SASP factors by combining data from the literature, and found 33% of them encoded by CSDE1 target transcripts (Figure S1F and Table S1).

A large number of transcripts were bound by CSDE1 in both proliferating and senescent conditions. To understand dynamic CSDE1 binding to these targets we conducted bioinformatics DESeq2 analysis to detect differential binding between EV and Ras samples. This analysis revealed 379 and 703 peaks with lower or higher density in senescence, respectively (Figure 3F and Table S2). Peaks with higher density (i.e. sites with higher binding to CSDE1) belonged to 459 genes enriched in pathways related to signaling and translation, while peaks with lower density belonged to 326 genes enriched in p53 signaling and cholesterol metabolism pathways. There was a minor overlap between the two groups (Figure S1G).

CSDE1 showed preferential binding to the coding sequence (CDS) and the 3’-UTR, and to a lesser extent to the 5’-UTR, indicating binding to mature mRNA, as previously described (Wurth et al., 2016, 2019) (Figure 3G). DREME analysis of iCLIP tags yielded enrichment of motifs with a conserved “AAG” core and small variations depending on condition (Figure 3H). Finally, RNA-maps analysis revealed a preference for CSDE1 positioning at the start codon and just upstream of the stop codon, consistent with previous observations and with a potential role of CSDE1 in translational regulation during senescence (Figure 3I).

### CSDE1 regulates the stability of SASP factor mRNAs

CSDE1 is known to regulate the stability and/or translation of its mRNA targets (Mihailovich et al., 2010). To identify CSDE1 targets changing at the steady state level during senescence, we performed RNA-Seq analysis of shCSDE1 vs shCTRL cells at early (7 dpi) and late (12 dpi) times upon senescence induction (Figure 4A). We performed 3 independent biological replicates per time point and condition (see PCA analysis in Figure S2A), and considered as significant those changes with padj ≤ 0.05 and L2FC ≥ 0.5. We obtained a total of 94 and 471 differentially expressed genes at 7 and 12 dpi, respectively, consistent with more pronounced remodeling of gene expression at later stages of senescence (Figure 4B and Table S3). Comparison with CSDE1 targets in senescent cells revealed a total of 24 targets regulated only at 7 dpi; 192 targets regulated only at 12 dpi and 22 targets regulated at both time points (Figure 4B). Downregulation upon CSDE1 depletion was predominant over upregulation, especially for iCLIP targets at early time points (Figure 4C and S2B), suggesting a role of CSDE1 in maintaining mRNA levels.

**Figure 4.**
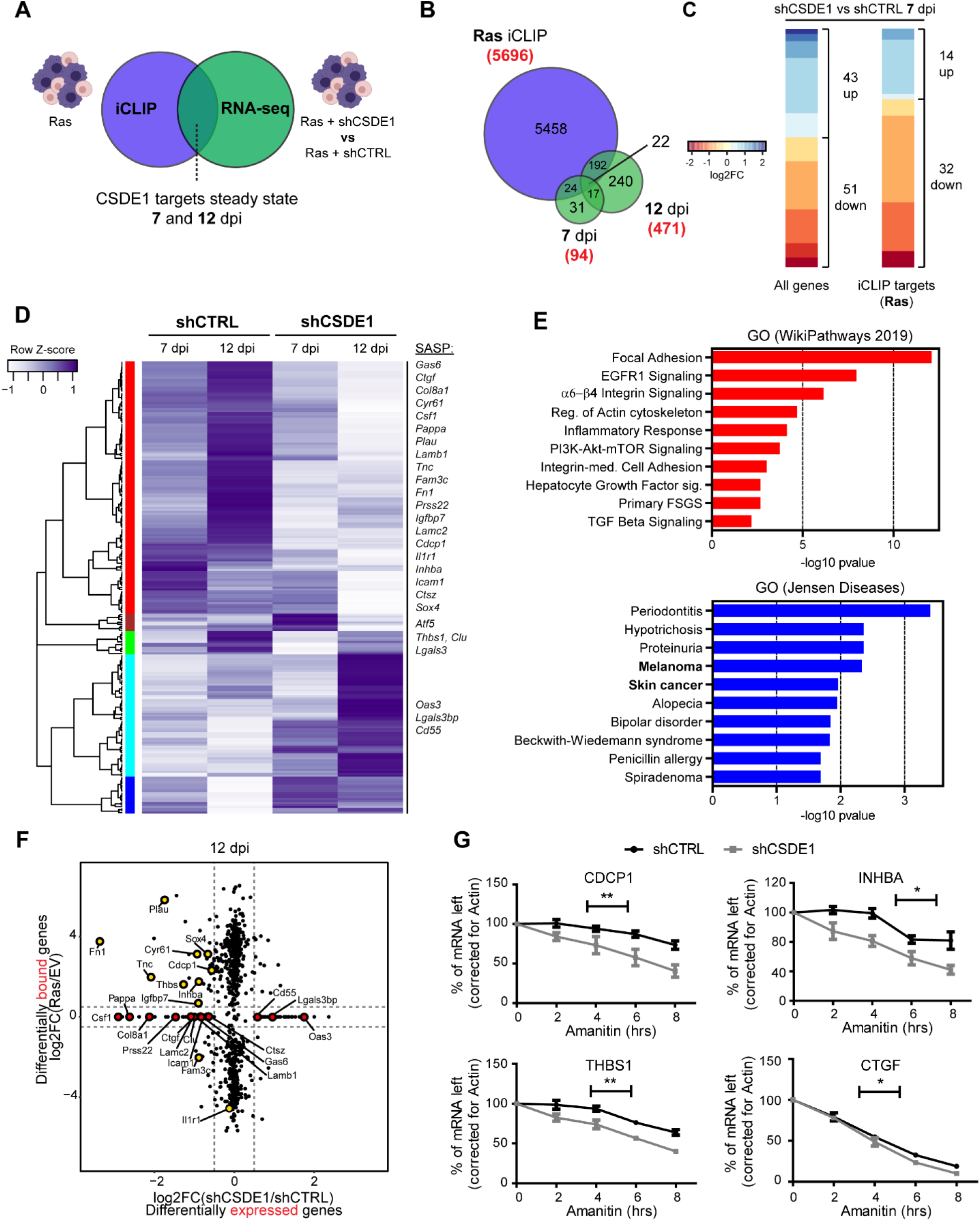
CSDE1 regulates the stability of SASP factor mRNAs. **(A)** Experimental strategy to identify CSDE1 targets regulated at the steady state level in senescent cells. **(B)** Overlap of CSDE1 targets with genes changing at 7 and 12 dpi after CSDE1 depletion. **(C)** Heatmaps of changes in the levels of all mRNAs or only CSDE1 targets at 7 dpi. **(D)** Unbiased hierarchical clustering of all changes in CSDE1 targets (238) observed at 7 and 12 dpi. Clusters are colored (left). SASP factor mRNAs are indicated. **(E)** GO analysis of the red and cyan/blue clusters. **(F)** Correlation of changes in mRNA binding and expression of CSDE1 targets (12 dpi). SASP factor transcripts are highlighted with a colored circle: yellow, differential binding; red, de novo binding. Note that the latter do not have a value in EV cells and, thereby, they are collapsed in the X-axis. **(G)** CSDE1 promotes the stability of SASP factor mRNAs. Transcription was inhibited with alpha-amanitin in shCTRL and shCSDE1 cells (12 dpi), and the levels of SASP factor transcripts followed at several times thereafter by RT-qPCR (n= 5). The data were corrected for actin and normalized to the initial time point. Linear regression analysis was conducted to test the significance of differences between mRNA decay slopes (*p-value <0.05, **p-value < 0.01). Error bars represent SEM.

Unbiased hierarchical clustering of all iCLIP targets regulated at the steady state level (238 genes) identified five main clusters (Figure 4D). The largest cluster (red) included transcripts with progressive downregulation in shCSDE1 cells, most of which showed progressive upregulation in senescent shCTRL cells. Strikingly, 20 out of 27 SASP transcripts bound and regulated by CSDE1 at these time points belong to this cluster, suggesting that CSDE1 promotes the SASP response by stabilizing SASP mRNAs. Changes in the levels of these transcripts upon CSDE1 depletion were validated by RT-qPCR in independent experiments, yielding a validation rate of 75% (Figure S2C). GO analysis of genes falling into this cluster showed enrichment in extracellular matrix remodeling and inflammation-related pathways, which fits well with a role of CSDE1 in SASP regulation (Figure 4E). Two other prominent clusters (cyan and blue) included transcripts that were downregulated during senescence and upregulated in shCSDE1 cells (Figure 4D). Interestingly, when we interrogated the Jensen Diseases database for genes in these two clusters, skin cancer and melanoma appeared among the most significantly related diseases (Figure 4E). We performed the same hierarchical clustering for all transcripts (CSDE1 targets or not) whose levels significantly changed in shCSDE1 cells. The results showed preferential inclusion of SASP transcripts among those downregulated upon CSDE1 depletion (Figure S2D). This shows that CSDE1 is required for SASP production via upregulation of SASP mRNA levels. We next wondered whether there was a correlation between dynamic CSDE1 binding to its targets and regulation of their levels. We interrogated the CSDE1 target group for differential binding and expression. Strikingly, all but one SASP factor transcript were differentially bound and regulated by CSDE1, with the vast majority showing increased binding and expression in senescent cells (Figure 4F; red, de novo binding; yellow, differential binding). These results suggested that CSDE1 stabilizes SASP factor mRNAs. To directly test this, we followed the levels of SASP factor mRNAs in shCTRL and shCSDE1 cells (12 dpi) after inhibiting transcription with alpha-amanitin. RT-qPCR at various points after inhibition indicated reduced stability upon CSDE1 depletion of 4 out of 6 SASP factor mRNAs tested (Figures 4G and S2E). Altogether, these data indicate that CSDE1 stabilizes SASP factor mRNA levels, consistent with a role of CSDE1 in promoting the SASP.

### Regulation of translation by CSDE1 during senescence

Our previous studies in melanoma revealed that CSDE1 promotes oncogenesis by regulating the translation of key EMT markers (Wurth et al., 2016, 2019). To understand whether translation regulation also contributes to the tumor-suppressive properties of CSDE1, we undertook polysome profiling followed by sequencing (PP-Seq) (Figure 5A). Given the limiting material of our primary cell system, we employed a previously reported non-linear sucrose gradient that concentrates highly translated mRNAs (> 3 associated ribosomes) in only 2 fractions, avoiding the loss of material typically associated to this laborious technique (Liang et al., 2018) (Figure 5B). No differences in the polysomal profile were detected upon CSDE1 depletion, indicating that CSDE1 did not cause defects in global translation (Figure 5C). To identify mRNAs regulated at the translation level by CSDE1, we performed PP-Seq in four independent biological replicates of CSDE1-depleted versus control cells at 12 dpi (Figure S3A). We applied poly-A RNA-Seq to both polysome and sub-polysomal (here referred to as ‘monosome’) fractions and analyzed our data with the anota2seq algorithm (Oertlin et al., 2019). Anota2seq identifies changes in polysome vs monosome association of mRNAs leading to three different regulatory outcomes: changes in mRNA abundance, changes in mRNA translation, and mRNA buffering, a process by which changes in mRNA abundance are counteracted at the translation level (or vice versa) in order to preserve protein levels (Atlasi et al., 2020; Blevins et al., 2019). In line with CSDE1 roles in translation regulation and mRNA stability, we found many genes regulated at both levels (Figure 5D). Interestingly, our analysis also revealed a large number of buffered genes (Figure 5D and Table S4), with CSDE1 targets in this category triplicating that of translationally regulated genes (Figure S3B). Curiously, GO analysis of both categories yielded cellular senescence as one of the most significantly enriched pathways only in the buffered gene category (Figure S3C). We reasoned that changes in translation of important senescence-related genes were being counteracted at the mRNA level (i.e. buffered) during these early times of senescence bypass and that, if relevant, these changes should be consolidated in immortalized cells. Thereby, we measured the polysome-to-monosome association of buffered CSDE1 iCLIP targets with described roles in senescence, including FOXO1, FOXO3, HIPK1, HIPK3, PTEN, SMAD3 and YBX1, in immortalized PMK. Four of the seven transcripts analyzed showed a significantly different association to polysomes in iPMK cells compared to control (Figure 5E). Of these, only YBX1 and PTEN showed changes consistent with senescence bypass, as translation of PTEN mRNA-encoding a tumor suppressor-decreased while translation of YBX1 mRNA-encoding a pro-oncogenic RBP recently shown to block replicative senescence (Kwon et al., 2018)-increased. Indeed, Western blot analysis confirmed a strong upregulation of YBX1 protein levels in iPMK, and already in CSDE1 depleted cells at 12 dpi (Figure 5F).

These data suggest that CSDE1-mediated translational regulation of senescence-associated genes, among them YBX1, might be essential for the establishment OIS.

**Figure 5.**
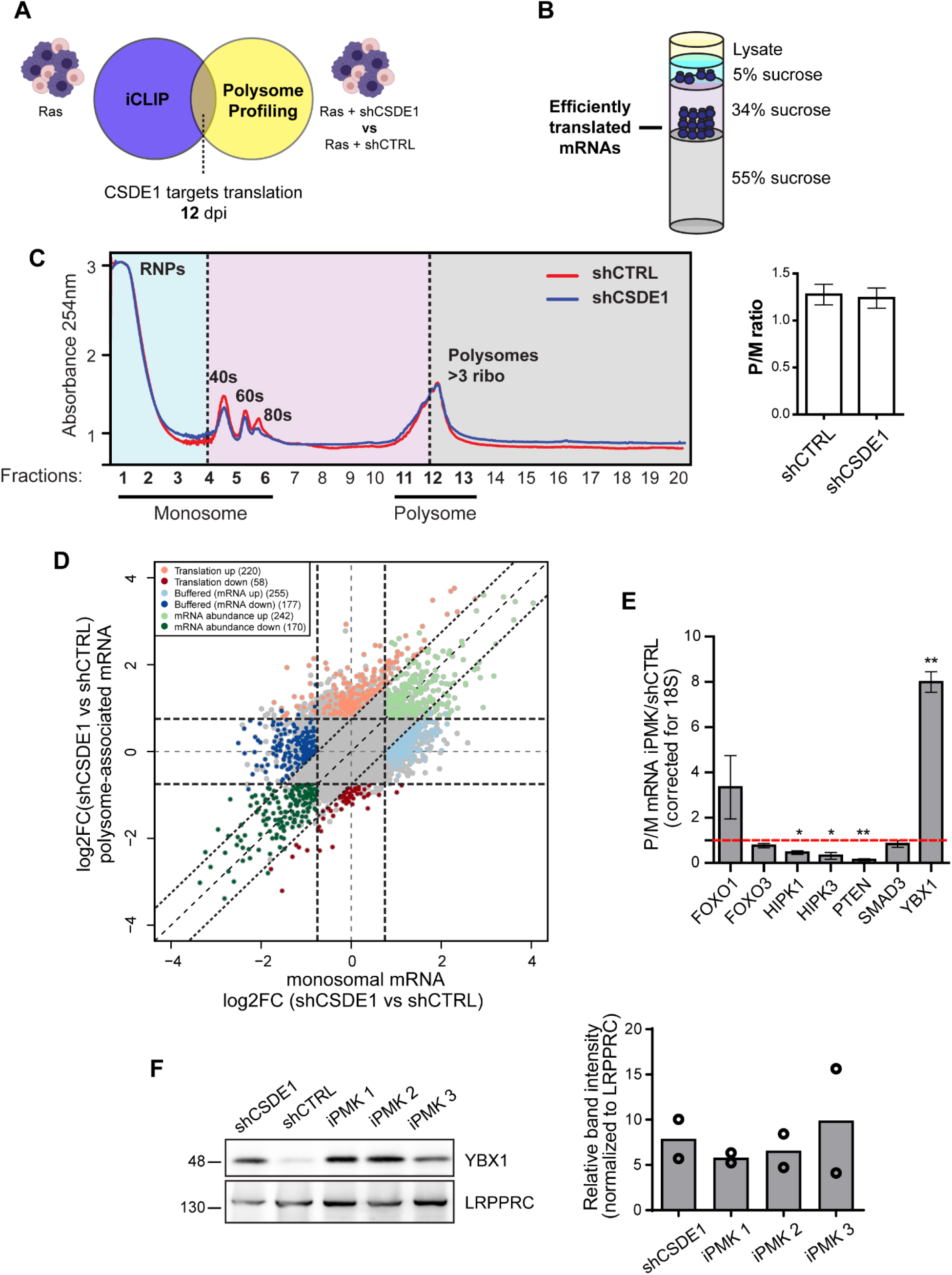
CSDE1 regulates the translation of YBX1. **(A)** Experimental strategy to identify CSDE1 targets regulated at the level of translation during senescence. **(B)** Schematic representation of the optimized non-linear gradient adopted for polysome profiling experiments. **(C)** Left, typical polysome profiles of shCTRL and shCSDE1 cells. The polysomal and sub-polysomal (herein referred to as ‘monosome’) fractions are indicated. Right, quantification of the polysome/monosome area ratio in shCTRL and shCSDE1 profiles (n= 3). **(D)** Scatter plot of polysome vs monosome-associated mRNA in shCSDE1 and shCTRL cells. Genes are colored according to their mode of regulation derived from anota2seq analysis (n= 4, FDR ≤ 0.2). **(E)** Translational changes are consolidated in immortal PMK. The polysomal distribution of selected buffered mRNAs was evaluated in iPMK and compared with shCTRL cells (n= 3). Significance was assessed by one sample t-test (*p-value <0.05, **p-value < 0.01). Error bars represent SEM. **(F)** YBX1 levels increase following CSDE1 depletion. Quantification from two independent experiments is shown on the right.

### YBX1 depletion from iPMK rescues the senescent phenotype

To assess the relevance of CSDE1-mediated translational regulation of YBX1 in OIS, we depleted YBX1 from iPMK and tested whether senescence was restored in these otherwise proliferating cells. We infected iPMK with a YBX1-directed shRNA and monitored the appearance of enlarged, flattened senescent cells for 17 days. Indeed, enlarged cells appeared in the three iPMK cell lines (Figure 6A). This effect was not due to recovery of CSDE1 expression (Figure 6B). BrdU incorporation assays confirmed reduced proliferation of iPMK cells after YBX1 depletion (Figure 6C). These data indicate that translational repression of YBX1 by CSDE1 is important to maintain OIS.

**Figure 6.**
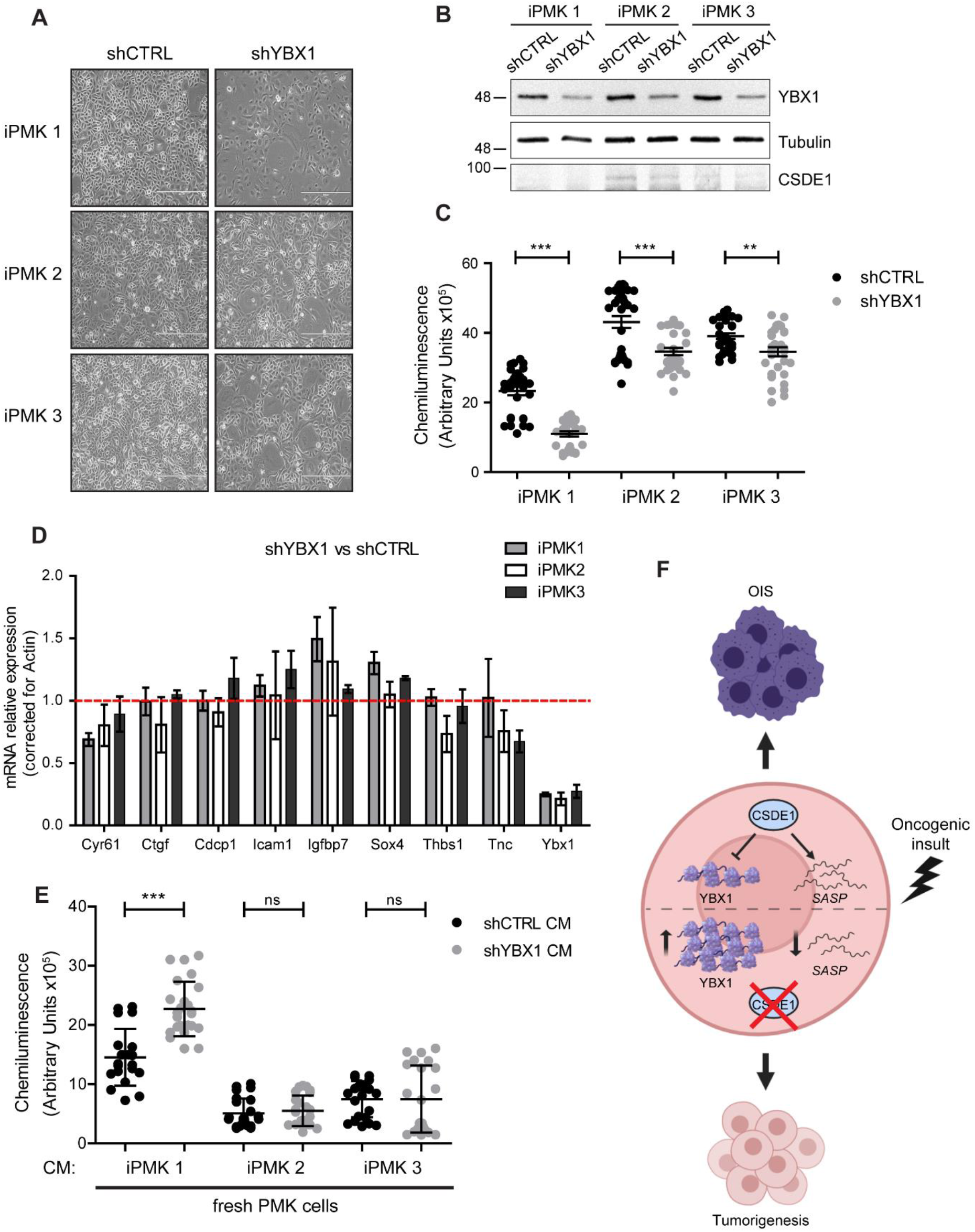
YBX1 depletion rescues the senescent phenotype. **(A)** Bright-field microscopy images of iPMK cells after 17 days of YBX1 depletion. **(B)** Analysis of YBX1 and CSDE1 levels by Western blot. **(C)** BrdU incorporation assays on shCTRL and shYBX1 iPMK at 10 days after depletion (n= 3, each with 10 technical replicates). Significance was assessed by unpaired Student’s t-test (**p-value < 0.01, ***p-value < 0.001). **(D)** SASP mRNA levels do not increase after YBX1 depletion. Transcript levels were assessed by RT-qPCR, corrected for actin and normalized to the levels in shCTRL cells (red line) (n= 9). Significance was assessed by one sample t-test. Error bars represent SEM. **(E)** BrdU incorporation assay of recipient PMK cells after 5 days of CM treatment (n= 3, each with 8 technical replicates). **(F)** Model for the role of CSDE1 in senescence. CSDE1 promotes senescence by following two independent mechanisms: stabilization of SASP factor mRNAs, and repression of YBX1 translation.

YBX1 has been previously shown to inhibit replicative senescence by repressing the translation of SASP factor mRNAs (Kwon et al., 2018). Comparison of the set of SASP factors regulated by YBX1 during replicative senescence with those regulated by CSDE1 during OIS revealed no overlap (data not shown). Nevertheless, to formally exclude the possibility that the effects of CSDE1 in SASP mRNA regulation are secondary to regulation of YBX1, we first analyzed SASP mRNA expression in iPMK cells. RNA-Seq indicated a profound remodeling of gene expression in iPMK compared to senescent shCTRL cells, probably fundamental for the immortalization and transformation process (Figure S4A and Table S5). A total of 56 SASP mRNAs showed reduced expression in iPMK, half of which are direct CSDE1 targets (Figure S4B). At an earlier time of senescence bypass (12 dpi), a lesser number of SASP factor mRNAs are downregulated, pointing to a progressive shut-down of the SASP signal (Figure S4C). Nine transcripts were kept low both at 12 dpi and in iPMK (Figure S4C and Table S5), suggesting that they are relevant SASP components regulated by CSDE1. Importantly, depletion of YBX1 did not result in upregulation of these transcripts in any iPMK cell line (Figure 6D). Furthermore, conditioned media experiments showed no decrease in the proliferation of recipient cells when incubated with media from shYBX1 cells (Figure 6E). These data indicate that CSDE1 regulation of SASP factor mRNA levels is independent of YBX1.

In summary, we show that CSDE1 follows a two-branched molecular mechanism to coordinately regulate cell proliferation and SASP expression during senescence: on one hand, CSDE1 stabilizes SASP factor mRNAs, and on the other it represses the translation of the pro-oncogenic RBP YBX1 (Figure 6F). These data place CSDE1 as an important post-transcriptional mediator of cellular responses against persistent oncogene signaling.

## DISCUSSION

Senescence is a cellular stress response mechanism associated with dramatic and dynamic changes in transcription (Hernandez-Segura et al., 2017). The role of post-transcriptional regulation in this process is, however, far from understood. To date, only a handful of stress-related RBPs have been involved in this process (Wang, 2012). Here we show that the RBP CSDE1 coordinates a post-transcriptional program to enforce proliferation arrest and SASP production during OIS.

Numerous lines of evidence indicate that the SASP and cell cycle are regulated by independent regulatory networks. For example, while cell cycle regulation depends on the cell cycle inhibitors p53 and p16^INK4a^, this is not the case for the SASP, which depends on a separate transcriptional program (Coppe et al., 2011; Kang et al., 2015). Accordingly, the SASP can be regulated independent of growth arrest (Nacarelli et al., 2019). CSDE1 behaves as a post-transcriptional orchestrator of both proliferative and SASP responses by using distinct mechanisms.

Proliferation arrest promoted by CSDE1 does not involve direct regulation of canonical cell cycle inhibitors. Rather, it involves translational repression of the pro-oncogenic factor YBX1, an RBP with functions in a broad range of gene expression processes, from transcription and splicing to mRNA stability and translation (Mordovkina et al., 2020). In cancer, YBX1 expression correlates with tumor growth and aggressiveness, consistent with its capacity to promote translation of EMT markers in breast cancer cells, or of HIF1α in high-risk sarcomas (El-Naggar et al., 2015; Evdokimova et al., 2009; Maurya et al., 2017). In senescence, YBX1 has been shown to repress the transcription of p16^INK4a^ in MEFs to oppose OIS (Kotake et al., 2013), and to prevent replicative senescence of primary human keratinocytes by promoting the translation of proliferation and self-renewal transcripts, while inhibiting the synthesis of cytokines-notably IL8 and CXCL1-that are part of the SASP (Kang et al., 2015; Kwon et al., 2018). In line with these reports, we find that YBX1 also counteracts OIS in primary mouse keratinocytes, although the mechanism is unrelated to regulation of IL8, CXCL1, or the SASP altogether, because depletion of YBX1 from immortalized keratinocytes reinstates senescence without restoring the SASP.

Independent of YBX1 regulation, CSDE1 promotes the SASP by stabilizing SASP factor mRNAs. Regulation of SASP factor mRNA stability is a recognized mechanism to control the SASP. The AU-rich element binding protein ZFP36L1 has been shown to play a role opposite to CSDE1. In human fibroblasts undergoing OIS, mTOR promotes the translation of MAPKAPK2, which in turn phosphorylates ZFP36L1 inhibiting its capacity to promote the degradation of SASP factor mRNAs (Herranz et al., 2015). Regulation of the stability of other mRNAs has also been reported to contribute to OIS modulation. FXR1 destabilizes p21^CIP1^ mRNA and stabilizes TERC mRNA to suppress senescence. In agreement, silencing of FXR1 in oral cancer cells triggers cellular senescence, similar to depletion of YBX1 in our system (Majumder et al., 2016). In addition, regulation of mRNA stability impinges on replicative senescence. HuR, one of the first RBPs shown to play a role in senescence, regulates the stability of several target mRNAs to maintain the replicative capacity of cells (Chang et al., 2010; Lee et al., 2018; Wang et al., 2001). CSDE1 now adds to this small but growing list of RBPs regulating transcript levels with consequences for senescence. Among the SASP factor mRNAs consistently down-regulated after CSDE1 depletion, we find a group coding for proteins of the extracellular matrix (CLU, CYR61, CTGF, CDCP1, ICAM1, IGFBP7, THBS1, TNC) whose remodeling fits well with the tumorigenic potential of iPMK cells. How downregulation of each SASP target precisely contributes to senescence bypass and acquisition of tumorigenic traits is an interesting question for future research.

In addition to post-transcriptional regulation of YBX-1 and SASP factor mRNAs, CSDE1 regulates many other transcripts in a direction that is consistent with reinforcement of senescence. Examples are the translational repression of the proliferation marker Ki67 or the translational stimulation of the tumor suppressor PTEN, which contribute to translational reprogramming during OIS. Thus, CSDE1 emerges as a multi-purpose post-transcriptional coordinator of OIS.

Depletion of CSDE1 in PMK challenged with Ras leads to senescence bypass measured by several independent assays. The widely used senescence-associated β-gal assay (SA-β-gal) was, however, not useful in our system. While non-infected PMK and shControl cells scored as expected (i.e. negative SA-β-gal staining for the former and positive for the latter), both shCSDE1 cells and immortal keratinocytes showed high levels of β-galactosidase activity (data not shown), suggesting that lysosomal activity remains high in cells that bypass senescence after depletion of CSDE1. Importantly, CSDE1 depletion not only leads to senescence bypass but to cell transformation, indicating that CSDE1 is a tumor suppressor in PMK. Although this role is consistent with loss-of-function mutations in neuroendocrine tumors (Fishbein et al., 2017), it contrasts with a wealth of evidence showing pro-oncogenic functions for CSDE1 (Fang et al., 2005; Liu et al., 2020; Martinez-Useros et al., 2019; Tian et al., 2018; Wurth et al., 2016). Thus, CSDE1 performs context-specific functions in cancer. Interestingly, CSDE1 binds a highly overlapping set of transcripts both in melanoma cells and PMK, including transcripts that are functionally relevant in both conditions. However, binding does not necessarily imply a regulatory outcome and, if any, the direction of the outcome might be different. For example, while CSDE1 binds PTEN mRNA in both PMK and melanoma cells, CSDE1 promotes the synthesis of PTEN in PMK, while it reduces the levels of PTEN mRNA in melanoma (Wurth et al., 2016 and this work). In principle, this could be due to different CSDE1 isoforms or post-translational modifications present in each system, which endow CSDE1 with capacity to interact with distinct effectors. Deciphering what makes CSDE1 a tumor suppressor or an oncoprotein will be the goal of future research.

## ACKNOWLEDGMENTS

We thank Bill Keyes and members of his lab Mekayla Storer, Birgit Ritschka and Alba Mas for expert advice and help at the initial stages of this project. We also thank Jernej Ule for sharing reagents and expertise on iCLIP, Ola Larsson for guidance on Anota2seq, and Bill Keyes and Juan Valcárcel for carefully reading the manuscript. We acknowledge the CRG Bioinformatics Unit, the Genomics Facility and the CRG/UPF FACS Unit for cell sorting and high-throughput sequencing and analysis. M. I-F. and A. C. were supported by FPI fellowships from the Spanish Ministry of Science and Innovation (MICINN). F.G. was supported by grants from MICINN (PGC2018-099697-B-I00 and BFU2015-68741), ‘La Caixa’ Foundation (ID 100010434) under the agreement LCF/PR/HR17/52150016, the Catalan Agency for Research and Universities (2017SGR534) and the Centre of Excellence Severo Ochoa.

## AUTHOR CONTRIBUTIONS

R. A., M. I-F., and A. C. performed the experiments, S. B. performed the bioinformatics iCLIP analyses, A. R. provided technical support with mice, and F.G. conceived and supervised the project. R. A. and F. G. wrote the manuscript and all authors provided comments.

## RESOURCE AVAILABILITY

### Lead Contact

Further information and requests for resources and reagents should be directed to and will be fulfilled by the Lead Contact, Fatima Gebauer.

### Materials Availability

All unique and stable reagents generated in this study are available from the Lead Contact with a completed Material Transfer Agreement.

### Data and Code Availability

All software used in this study is listed in the Key Resources Table. Data generated in this study are available through Gene Expression Omnibus (GEO) under accession number GEO: GSE160002. Experimental model was created using graphics from https://www.biorender.com.

## EXPERIMENTAL MODEL AND SUBJECT DETAILS

### Mouse care

Wild-type C57BL/6J and Swiss Nude mice were housed in accordance with the Ethical Committee for Animal Experimentation (CEEA) of the Government of Catalonia.

### Tissue culture

Primary Mouse Keratinocytes (PMK) were obtained from newborn C57BL/6J mice (1-2 days old). Briefly, pups were sacrificed with an intraperitoneal injection of Duolethal (20 mg/ml), submerged in betadine, rinsed with PBS, washed with 70% ethanol, dried with paper and kept in PBS at 4°C. Skins were dissected and placed completely flat with dermis side down in a 100 mm tissue culture dish. For epidermis-dermis separation, the skin was incubated with 10 ml of Dispase II (2.5 mg/mL, Roche) at 4°C overnight. Within a laminar-flow hood, skins were taken and placed in a new 100 mm lid with the dermis side up while separating the epidermis with forceps, gently gilding away the dermis and discarding it. The epidermis was chopped using scissors and a blade until homogenous pieces were small enough to be collected with a 10 ml pipette. Skins were incubated under stirring with EMEM complete media during 40 min at room temperature. Using a cell strainer of 70 μm (Corning, 352350) the cell suspension was filtered into a 50 mL Falcon tube and centrifuged. Cell pellets were resuspended in the appropriate volume of EMEM complete media (Lonza, Cat 06-174G) supplemented with 8% of chelated FBS (Fetal Bovine Serum, see below), 0.05 mM CaCl2, 10 ng/ml EGF (Sigma-Aldrich) and 1% penicillin/streptomycin (Life technologies), and seeded in collagen-treated plates. To treat plates with collagen, Collagen I from rat tail (Invitrogen) was diluted to 1/60 in PBS and added to plates for at least 40 min at 37°C. Cells were incubated for 24 hours at 37°C in a CO2 incubator, and washed several times with PBS before adding CnT-Prime media (CELLnTEC).

For chelated FBS preparation, 500 ml of FBS (Invitrogen) were heat inactivated for 30 min at 56 °C and chelated with 150 gr of treated Chelex-100 Resin (BioRad) for 1 hr at room temperature with stirring. To treat the Chelex-100 resin, 150 gr of resin were added to 2 L of milliQ water and gently mixed by stirring overnight at room temperature. The following day, pH was adjusted to 7-7.5, the water decanted and the resin recovered with the help of 3MM paper. The chelated FBS was filtered with 0.45 μm filter and stored at −20°C.

## METHOD DETAILS

### Constructs

shCSDE1 and shCTRL hairpins were cloned into the XhoI and EcoRI sites of the pLMP retroviral vector. The pLMP vector contains a puromycin-IRES-GFP cistron (PIG) for both drug and FACS selection. shRNA inserts of 97-mer were identical to those used in Wurth et al., 2016. shYBX1 (ULTRA-3247270) and shCTRL in pLMP-d-mAmetrine vector (shERWOOD UltramiR) were purchased from transOMIC. The pLMP-d-mAmetrine vector contains a mAmetrine fluorescent protein under the control of the PGK promoter for FACS selection. MSCV-Empty vector (EV) was used as control for retroviral infection-related effects. H-Ras^V12^-expressing vector (Ras), containing the human H-RAS^V12^ gene in a pWZL backbone, was used to induce senescence. A single nucleotide change (G to C) leads to the oncogenic missense variant Gly12Val. Both the Ras and MSCV vectors contain an hygromycin resistance gene under the control of the PGK promoter for drug selection.

### Viral transduction

To generate retroviral particles, the 293T-based Phoenix ecotropic (E) packaging cell line (generated by Garry Nolan laboratory, Stanford University) was transfected with the plasmid of interest (EV, RAS, shCTRL, shCSDE1, shCTRL [expressing Ametrine], shYBX1). Prior to Phoenix E transfection, chloroquine (Sigma-Aldrich) was added to the media to a final concentration of 25 μM. Ten μg of the plasmid and 60 μl of 2.5 M CaCl2 were mixed with water up to 500 μl. After 10 min, 500 ul of 2X concentrated HEPES Buffered Saline (281 mM NaCl, 100 mM HEPES and 1.5 mM Na2HPO4 in water, pH 7.12) was added into the DNA mix dropwise while vortexing. The mixture was incubated 15 min at room temperature and then added dropwise to the cells. Medium was replaced 7 hours after transfection. The supernatant, containing viral particles, was collected after 48 hours and used to infect PMK. PMK were infected 48 hours after seeding, following three rounds of 2 hours incubation with 8 mL of virus-containing media. In the case of double infection, both supernatants were added simultaneously to PMK. Polybrene (Sigma-Aldrich) was added to the viral supernatant at 10 μg/ml to enhance retroviral infection. Cells were split 24 hours after the last infection, selected with of puromycin (1 μg/mL) for 48 hours and, in case of double infections, further selected with hygromycin (25 μg/mL). Cells were maintained in CnT-Prime media prior to assays.

### Fluorescence Activated Cell Sorting (FACS)

For GFP and mAmetrine sorting, cells were collected at 7 dpi and resuspended in CnT-Prime media. DAPI was added to a final concentration of 1 μg/mL in order to discard dead cells.

### Mouse xenografts

One million cells in 200 μl of CnT-Prime media and 200 ul of Growth Factor Reduced (GFR) Basement Membrane Matrix (Corning) were injected subcutaneously, bilaterally into the flanks of 8 week-old swiss nude mice (Charles River laboratory). Three mice were used per iPMK cell line. After 20-30 days, tumors were resected and weighted.

### Conditioned media

Conditioned media was collected at 7 dpi from shCTRL and shCSDE1 PMK, and 7 days after depletion from shCTRL and shYBX1 iPMK, filtered through a 0.45 μm filter, aliquoted and stored at −20°C. Conditioned media was then used to grow freshly isolated keratinocytes 24 hours after seeding. Conditioned media was replenished every 24 hours for 6 days.

### Colony Forming Assay

shCTRL and shCSDE1 PMK (7 dpi) were layered on J2P mitotically inactivated cells (a gift from Bill Keyes). Experiments were performed in duplicates at three different seeding densities in 6-well-plates: 500, 1000 and 2000 cells per well. Experiments were performed as described in (Jensen et al., 2010). Colonies were visualized with 1% Rhodamine B.

### BrdU incorporation assay

For BrdU experiments 20000 cells were seeded in collagen-treated 96-well-plates (Corning). BrdU was added for 18 hours to a final concentration of 100 μM. BrdU incorporation was measured using the Cell Proliferation ELISA kit (Roche #11-669-915-001) following the manufacturer’s instructions.

### RNA isolation and RT-qPCR

For RT-qPCR, RNA was extracted with either TRIzol reagent (Invitrogen) or Maxwell kit (Promega). Following RNA extraction, samples were subjected to DNase digestion using the Turbo DNA-free kit (Invitrogen). RNA was reverse-transcribed using 2.5 μM oligo(dT), 2.5 ng/μl random primers, 0.05 mM of a mix of dNTPs, 1 mM DTT, 1X first strand buffer, 1 μl of RNase OUT (Invitrogen), and 0.5 ul of SuperScript II (50 U/ul) (Invitrogen) in a final volume of 20 μl. The resulting cDNAs were diluted ¼ and used as templates for qPCR using the Power SYBR Green Master Mix (Applied Biosystems) following the manufacturer’s instructions.

### Immunoblotting

Protein extraction was performed with RIPA buffer (150 mM NaCl, 10 mM Tris-HCl pH 7.5, 0.1% SDS, 1% DOC, 5 mM EDTA, 1% Triton X-100 and 1X protease inhibitors) allowing for both nuclear and cytoplasmic protein extraction. Protein extracts were separated by SDS-PAGE, transferred to PVDF membranes, blocked 1 hour at room temperature with 5% low-fat milk in TBS-T (10 mM Tris-HCl pH 7.5, 100 mM NaCl, 0.1% Tween-20) and incubated with primary antibody overnight at 4°C. Membranes were then washed (3 times for 10 min) with TBS-T and incubated with Horseradish Peroxidase (HRP)-coupled secondary antibody for 1 hour at room temperature. After two washes of 10 min, proteins were detected by ECL (Perkin Elmer). Antibodies used are listed in the key Resource Table.

### iCLIP-Seq

Not-infected (NI), Empty Vector-infected (EV) and H-Ras^V12^-infected (Ras) PMK were cross-linked on ice at 0.15 J /cm^2^ with UV light at 254 nM. Immediately after irradiation, cells were lysed in 1 mL lysis buffer (50 mM Tris-HCl, pH 7.4; 100 mM NaCl; 1% NP-40; 0.1% SDS; 0.5% sodium deoxycholate). Cell extracts were sonicated using a bioruptor (Digenode) for 10 cycles of 30 seconds, level L at 4°C, and then cleared by centrifugation at 10000 rpm for 10 min at 4°C. RNA was then partially digested by adding 10 μl of 1:200 dilution of RNase I (Invitrogen), as well as 2 μl of Turbo DNase (Invitrogen) and incubating for 3 min at 37°C under shaking at 1100 rpm. CSDE1 protein was then captured by incubation with an affinity purified CSDE1 antibody conjugated to protein A Dynabeads (Invitrogen) for 4 hours at 4°C with gentle rotation. Beads were then washed 2x with 900 μl high-salt buffer (50 mM Tris-HCl pH 7.4, 1 M NaCl, 1 mM EDTA, 1% NP-40, 0.1% SDS, 0.5% sodium deoxycholate), 1x with 900 μl PNK wash buffer (20 mM Tris-HCl pH 7.4, 10 mM MgCl2, 0.2% Tween-20) and resuspended in 40 μl of PNK mix (30 μl water, 8 μl 5x PNK pH 6.5 buffer [350 mM Tris-HCl pH 6.5, 50 mM MgCl_2_, 5 mM DTT], 1 μl PNK enzyme [NEB], 0.5 μl RNasin [Promega], 0.5 μl of FastAP alkaline phosphatase [Thermo Fisher Scientific]) and incubated 40 min at 37°C under gentle shaking at 1100 rpm. Beads were washed 1x with 900 μl of ligation buffer (50 mM Tris-HCl pH 7.5, 10 mM MgCl2) and resuspended in 25 μl of ligation mix (6.8 μl water, 2.5 μl 10X ligation buffer, 0.8 μl 100% DMSO, 2.5 μl T4 RNA ligase I – high concentration [NEB], 0.4 μl RNasin [Promega], 0.5 μl PNK enzyme, 2.5 μl pre-adenylated adaptor L3-App [1 μM] conjugated to an infrared-dye [IRdye-800CW-DBCO (LICOR)], 9 μl 50% PEG8000). Ligation was performed at room temperature for 75 min flicking the tubes every 10 min. Beads were washed 2x with high-salt buffer, 1x with PNK wash buffer, resuspended in 20 μl of loading buffer and run on a 4-12% NuPAGE Bis-Tris gel (Invitrogen) in MOPS buffer according to the manufacturer's instructions. Protein-RNA complexes were transferred to a nitrocellulose membrane. Protein-infrared-RNA complexes were visualized with an Odissey machine setting of1.5 intensity for both 700 nm and 800 nm channel. RNA-protein complexes were isolated and treated with 200 μl PK buffer (10 mM Tris-HCl pH 7.4, 100 mM NaCl, 1 mM EDTA, 0.2% SDS) and 10 μl proteinase K (Roche) for 1 hour at 1100 rpm and 50°C. Solution was collected and RNA was isolated by phenol:chloroform extraction and ethanol precipitation over night at −20°C. The RNA pellet was resuspended in 7 μl RNA/primer mix (5.5 μl water, 1 μl irCLIP_ddRT_## primer,0.5 μl dNTP mix [10mM]) and incubated at 65°C for 5 min. RNA was reverse-transcribed by addition of 3 μl of RT mix (2 μl 5x SSIV buffer [Invitrogen], 0.5 μl 0.1 M DTT, 0.25 μl Superscript IV reverse transcriptase [Invitrogen], 0.25 μl of RNasin) and incubation for 5 min at 25°C, 5 min at 50°C and 5 min at 55°C. cDNA was gel-size selected by running samples on a precast 6% TBE-urea gel (Invitrogen) at 180V for 40 min and isolating the region between 75 and 200 nt. Gel slices were submerged and crushed in TE buffer, incubated for 1 hour at 37°C and 1100 rpm, flash frozen on dry ice, and incubated again for 1 hour at 37°C and 1100 rpm. cDNA was isolated by phenol:chloroform extraction and ethanol precipitation overnight at −20°C. cDNA was resuspended in 7 μl of water and then 8 μl of CircLigase II mix (5 μl water, 1.5 μl 10x CircLigase Buffer II, 0.75 μl 50 mM MnCl2, 0.75 μl Circligase II [Lucigen]) were added. Samples were incubated for 1 hour at 60°C for circularization and cDNA was isolated by phenol:chloroform extraction and ethanol precipitation overnight at −20°C. Pellets were resuspended in 10 μl of water. Amplification was performed using 4 μl cDNA, 1 μl primer mix P5/P3 solexa (10 μM each); 20 μl Phusion HF Master mix [NEB], 15 μl water and the following PCR programme: 98°C for 40 sec, [98°C for 20 sec, 65°C for 30 sec, 72°C for 45 sec]x18 cycles, 72°C for 3 min, 4°C. Libraries were loaded on a precast 6% TBE gel (Invitrogen) and run at 180V for 30 min. Gel was stained with Sybergreen I (Invitrogen) and the region between 150-300 nt was excised. Gel slices were submerged in Crush-Soak Gel buffer (500 mM NaCl, 1 mM EDTA, 0.05% SDS) and incubated at 65°C for 2 hours with thermomixer settings of 15 sec at 1000 rpm, 45 sec rest. cDNA was isolated by phenol:chloroform extraction and ethanol precipitation over night at −20°C and resuspended in 10 μl of water. Libraries were sequenced using Illumina HiSeq 2500 sequencer (50 bp single-end reads) at the CRG Genomics Facility. Data were analyzed as described in computational methods.

### RNA-Seq

For total RNA-Seq, 500 ng of total RNA were depleted of ribosomal RNA using the Ribo-Zero rRNA removal kit (Epicentre). The resulting RNA was then used for library preparation using the TruSeq small RNA Sample Prep Kit (Illumina) according to the manufacturer's protocol. After purification on gel and amplification, final libraries were analyzed using Agilent DNA 1000 chip to estimate the quantity and check the size distribution, and were then quantified by qPCR using the KAPA Library Quantification Kit (Kapa Biosystems) prior to amplification with Illumina’s cBot. Libraries were sequenced using a HiSeq 2500 sequencer (50 bp single-end reads) at the CRG Genomics Facility. Data were analyzed as described in computational methods.

### Polysome Profiling-Seq

Polysome Profiling was adapted from Liang S. et al, 2017. Media was discarded from shCTRL and shCSDE1 cells (12 dpi – one 100 mm plate per condition), plates were placed on ice and immediately washed with an ice-cold solution of 1x phosphate-buffered saline (PBS) and cycloheximide (100 μg/mL, Sigma-Aldrich). Cells were then lysed in 200 μl hypotonic lysis buffer (5 mM Tris–HCl pH 7.5, 2.5 mM MgCl_2_, 1.5 mM KCl, 100 ug/mL cycloheximide, 2 mM DTT, 0.5% Triton, 0.5% sodium deoxycholate) and left on ice for 10 min. Extracts were cleared by centrifugation at 12’274 rpm for 7 min and the OD was measured (between 2.5 and 4 OD per experiment were loaded onto the gradient). Samples were brought to a final volume of 300 μl if necessary with lysis buffer and loaded on a non-linear sucrose gradient prepared by layering different percentage of sucrose solutions (5%, 34% and 55%) into an Open-Top Thinwall Ultra-Clear Tube (Beckman, 344059). A 5x gradient buffer (100 mM HEPES pH 7.6, 500 mM KCl, 25 mM MgCl_2_) was used to prepare sucrose solutions (1x final concentration of gradient buffer [v/v] adjusted with water). The samples were centrifuged at 36000 rpm for 2 hours at 4°C in a SW 41 Ti rotor and a Beckman Coulter Ultracentrifuge Optima L-90K. Samples were eluted using the gradient station (BioComp) and fractions (1-6 for monosomal RNA and 11 and 12 for polysomal RNA) were collected with Gradient Fractionator (BioComp) coupled with a fraction collector (Gilson). The precise location of the fractions along the UV-tracing was monitored using the gradient profiler v1.25 (BioComp) software. TRIzol reagent (Invitrogen) was immediately added to each fraction and the RNA was extracted according to the protocol of the manufacturer. For PolyA RNA-Seq, libraries were prepared using the TruSeq Stranded mRNA Sample Prep Kit v2 (Illumina) according to the manufacturer's protocol. Libraries were sequenced using a HiSeq 2500 sequencer (50 bp single-end reads) at the CRG Genomics Facility. Data were analyzed as described in computational methods.

### RNA-immunoprecipitation (RIP)

EV PMK cells cultured in 100 mm plates were washed with PBS and lysed in 200 μl lysis buffer (50 mM HEPES pH 7.4, 150 mM NaCl, 1 mM MgCl_2_, 2 mM EDTA, 1% NP-40, 1 mM DTT, 100 U/mL RNasin [Promega], 1X protease inhibitor) at 7 dpi. The suspension was incubated for 10 min on ice and centrifuged for 10 min at 14000 rpm at 4°C. Six hundred μg of the supernatant were incubated for 4 hours under rotation at 4°C with 20 μl of protein A Dynabeads (Invitrogen) containing anti-UNR antibody or rabbit IgG as control in a final volume of 600 μl. Ten percent of this suspension (60 μl) were taken for input RNA extraction. Beads were prepared as follows: 20 μl of beads were washed three times with 10 volumes IPP500 buffer (20 mM HEPES pH 7.9, 500 mM NaCl, 1.5 mM MgCl_2_, 0.5 mM DTT, 0.05% NP40) and subsequently bound to 6 μg of anti-UNR antibody or to rabbit IgG for 1 hour at room temperature. Beads were then washed twice with 10 volumes of IPP500 buffer and twice with 10 volumes of IPP150 buffer (20 mM HEPES pH 7.9, 150 mM NaCl, 1.5 mM MgCl_2_, 0.5 mM DTT, 0.05% NP-40) to remove excess antibody. Following incubation with the extract, beads were washed 4 times with 5 volumes of IPP150 buffer. Then, 200 μl of TRIzol were added to both beads and inputs and RNA was recovered following the manufacturer’s instructions. The RNA was then reverse transcribed and analyzed by RT-qPCR. The amount of precipitated RNA was normalized to the amount of the RNA in the input (arbitrary units).

### RNA half-life measurement

shCTRL and shCSDE1 cells were GFP-sorted at 7 dpi and plated in a six-well plate. Cells were treated with 5 μg/mL of alpha-amanitin (Sigma) and total RNA was extracted at time-points 0, 2, 4, 6, and 8 hours post-incubation. RNA levels were analyzed by RT-qPCR as described above using gene-specific oligonucleotides indicated in Table S6. Targets RNA levels at different time points were corrected for Actin RNA levels and normalized to the initial time point.

## QUANTIFICATION AND STATISTICAL ANALYSIS

Statistical significance in this study was determined by one-sample t-test analysis or unpaired, two-tailed Student’s t test with unequal variance at *p < 0.05, **p < 0.01, and ***p < 0.001 as specified in the legends of each figure. Error bars represent the standard error of the mean (SEM).

All statistical analyses were performed and graphics were generated using GraphPad Prism 5 software.

## COMPUTATIONAL METHODS

### iCLIP-Seq

#### iMaps analysis

iCLIP data were processed following the iCount Primary Analysis (Consensus mapping) pipeline from the iMaps website (https://imaps.genialis.com/iclip). Briefly, fastq files were demultiplexed and Cutadapt was used to trim off adapter sequences. Reads were first mapped to tRNA/rRNA reference. Unmapped reads where then mapped to the mouse genome (GRCm38/mm10) using the STAR aligner with default options. iCount xlsites was used to determine the number of unique crosslinked sites (unique cDNA molecules) for any given position with default options. iCount peaks was used to find positions with high density of cross-linked sites. Only peaks showing an FDR < 0.05 were considered significant and taken for subsequent steps. iCount RNA-maps was used to examine all cross-links that are located in vicinity of genomic landmarks.

Below, summary of sequencing reads:

|  | Total input reads | After Adapter Trimming | After rRNA/tRNA filtering | Uniquely mapped | After PCR dup. removal |
| --- | --- | --- | --- | --- | --- |
| NI_1 | 34527262 | 34025581 | 28162011 | 19081424 | 5704970 |
| NI_2 | 54797510 | 54107608 | 39384652 | 25988310 | 7519380 |
| NI_3 | 26626994 | 26444912 | 19532144 | 12956531 | 2435930 |
| NI_1_noUV | 128582 | 109108 | 72667 | 11390 | 4549 |
| NI_2_noUV | 2993958 | 2432656 | 1842479 | 232899 | 78402 |
| NI_3_noUV | 484438 | 332814 | 292913 | 34572 | 15769 |
| EV_1 | 14804464 | 13870767 | 11231980 | 6292141 | 639016 |
| EV_2 | 28744964 | 25691800 | 19622194 | 10440270 | 948712 |
| EV_3 | 37398668 | 35004134 | 27555556 | 15231900 | 5866820 |
| EV_1_noUV | 3300931 | 2982251 | 1162541 | 265889 | 27255 |
| EV_2_noUV | 1086849 | 945719 | 523305 | 80442 | 12603 |
| EV_3_noUV | 230486 | 205776 | 123035 | 35065 | 13691 |
| Ras_1 | 43114101 | 42475449 | 34175154 | 22565303 | 8509070 |
| Ras_2 | 74668905 | 73804303 | 59651086 | 39216341 | 14951200 |
| Ras_3 | 57890083 | 56879193 | 45002004 | 28511281 | 10421800 |
| Ras_1_noUV | 856312 | 782310 | 452851 | 65112 | 38491 |
| Ras_2_noUV | 4666762 | 4101968 | 1995249 | 281011 | 132791 |
| Ras_3_noUV | 1994226 | 1761559 | 1059612 | 159576 | 84915 |

### Differentially binding analysis

The iMAP crosslinked sites filtered for FDR < 0.05 were used. Ras BAM files were down-sized so as to match the coverage of EV BAM files (about 1 million reads). The down-sampling was done with samtools view (Li et al., 2009) with parameter “-s” set to 0.15 (H-ras replicate 1), 0.08 (H-ras replicate 2) and 0.12 (H-ras replicate 3). The down-sized Ras samples were then used for the next steps of the differential binding analysis. Raw counts over the iMAP crosslinked sites were calculated using the featureCounts function implemented in the Rsubread (v2.0.1) R/Bioconductor package (Liao et al., 2019) with options: isPairedEnd=ispaired, allowMultiOverlap=FALSE, useMetaFeatures=TRUE, countMultiMappingReads=FALSE, strandSpecific=1 (for iCLIP data: forward stranded) or strandSpecific=2 (for RNA-seq data: reverse stranded) DESeq2 version 1.26.0 (Love et al., 2014) was used to assess differentially bound peaks, as described by the DESeq2 main developer in: https://support.bioconductor.org/p/61509/.

The design used was as follow:

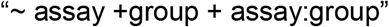

assay being the type of experiment (RNA-seq/iCLIP), group being the experimental groups (EV, Ras). The interaction term “assay:group” represents the ratio of ratios, for example, in the case of comparing Ras vs EV: (iCLIP for Ras /RNA-seq for Ras) / (iCLIP for EV/RNA-seq of EV). The likelihood ratio test (LRT) is then used to remove the interaction term in the reduced model, which allows to calculate the differential binding of iCLIP peaks, removing any artifact the RNA-seq experiment could introduce.

DESeq2 code:

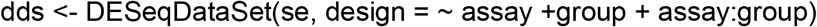

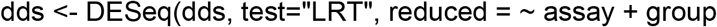

### Binding motif analysis

DREME v5.0.2 (Bailey, 2011) was used to find de novo motifs around the crosslinked regions. Motifs were searched +/− 10bp around each crosslinked region. DREME was run with parameters “-dna -norc -g 200 -mink 4 -maxk 7”.

### UCSC tracks

Deeptools (Ramirez et al., 2016) bigwigMerge was used to merge the 3 iCLIP replicates of each experimental condition together, and the 3 replicates of each RNA-seq experimental condition together. Deeptools bigwigCompare was then used to subtract the RNA-seq abundance from the iCLIP signal

### RNA-Seq

Quality of the fastq files was assessed with FastQC version 0.11.5 (Andrews, 2010). Reads were aligned with STAR version 2.5.3a (Dobin et al., 2013) to the mouse genome (release 88 of the ENSEMBL annotation) (Cunningham et al., 2019), using the “--quantMode GeneCounts” option for counting the number of reads aligning to each gene (the counts corresponding to the “reverse” strand were kept).

The R/Bioconductor (Huber et al., 2015; R Core, 2019) “DESeq2” (version 1.26.0) (Love et al., 2014) package was used to calculate the pairwise differential expression of genes between experimental groups.

The PCA (Principal Component Analysis) was computed with the “prcomp” function from the “stats” package, and plotted with the “ggplot2” package (Wickham, 2016). Hierarchical clustering was performed using an Rplatform and a heatmap described as using a function of heatmap.2 in gplots version 2.17.0 (Warnes, 2015).

Below, summary of sequencing reads:

|  | Total input reads | Uniquely mapped |
| --- | --- | --- |
| shCTRL_1_7dpi | 22227169 | 18936104 |
| shCSDE1_1_7dpi | 23295710 | 19774483 |
| shCTRL_2_7dpi | 21956045 | 18742114 |
| shCSDE1_2_7dpi | 25321613 | 21706799 |
| shCTRL_3_7dpi | 24630415 | 20794131 |
| shCSDE1_3_7dpi | 26045194 | 21623340 |
| shCTRL_1_12dpi | 26014075 | 21339896 |
| shCSDE1_1_12dpi | 23295875 | 19597477 |
| shCTRL_2_12dpi | 22558805 | 19280798 |
| shCSDE1_2_12dpi | 21327257 | 18024534 |
| shCTRL_3_12dpi | 19724662 | 16766499 |
| shCSDE1_3_12dpi | 23751094 | 20056924 |
| iPMK_1 | 19203370 | 16613018 |
| iPMK_2 | 19141924 | 16580368 |
| iPMK_3 | 18574214 | 15993674 |

### Polysome Profiling-Seq

Quality of the fastq files was assessed with FastQC version 0.11.5 (Andrews, 2010). Reads were aligned with STAR version 2.5.3a (Dobin et al., 2013) to the mouse genome (release 88 of the ENSEMBL annotation) (Cunningham et al., 2019), using the “--quantMode GeneCounts” option for counting the number of reads aligning to each gene (the counts corresponding to the “reverse” strand were kept). Anota2seq (Oertlin et al., 2019) was used to classify transcripts into three modes for regulation of gene expression: changes in mRNA abundance, changes in translational, and translational buffering. Counts from genes which had at least one read in each sample were scaled using TMM normalized library sizes (Robinson and Oshlack, 2010). Log2 counts per million were then computed using the voom function of the limma R package (Ritchie et al., 2015) as implemented in anota2seq.The following significance thresholds were used in the anota2seqSelSigGenes function: minEff = log2(0.75); maxPAdj = 0.2 (FDR < 0.2), selDeltaPT = selDeltaTP = selDeltaP = selDeltaT = log2(0.75); maxSlopeTranslation = 2; minSlopeTranslation = -1; maxSlopeBuffering = -2; minSlopeBuffering = 1. The different regulatory modes are visualized in scatter plots of polysome-associated mRNA log2Fold Change vs monosome-associated mRNA log2Fold Change (shCSDE1 vs shCTRL).

The PCA (Principal Component Analysis) was computed with the “prcomp” function from the “stats” package, and plotted with the “ggplot2” package (Wickham, 2016).

Below, summary of sequencing reads:

|  | Total input reads | Uniquely mapped |
| --- | --- | --- |
| shCTRL_mono_1 | 21082376 | 12511014 |
| shCSDE1_mono_1 | 20837197 | 12362756 |
| shCTRL_mono_2 | 24093474 | 11473491 |
| shCSDE1_mono_2 | 16571824 | 9683935 |
| shCTRL_mono_3 | 19416009 | 10372668 |
| shCSDE1_mono_3 | 18208506 | 10222359 |
| shCTRL_mono_4 | 19365479 | 10845757 |
| shCSDE1_mono_4 | 19896230 | 12242894 |
| shCTRL_poly_1 | 24048479 | 18770818 |
| shCSDE1_poly_1 | 25514411 | 19951591 |
| shCTRL_poly_2 | 23747355 | 17938067 |
| shCSDE1_poly_2 | 23090812 | 17688608 |
| shCTRL_poly_3 | 23967215 | 19659930 |
| shCSDE1_poly_3 | 19289250 | 15230830 |
| shCTRL_poly_4 | 21062658 | 16517680 |
| shCSDE1_poly_4 | 20324698 | 15759101 |

### Gene ontology

For all gene ontology analyses the enrichR webtools was used (Kuleshov et al., 2016).

## SUPPLEMENTAL FIGURE LEGENDS

**Figure S1.**
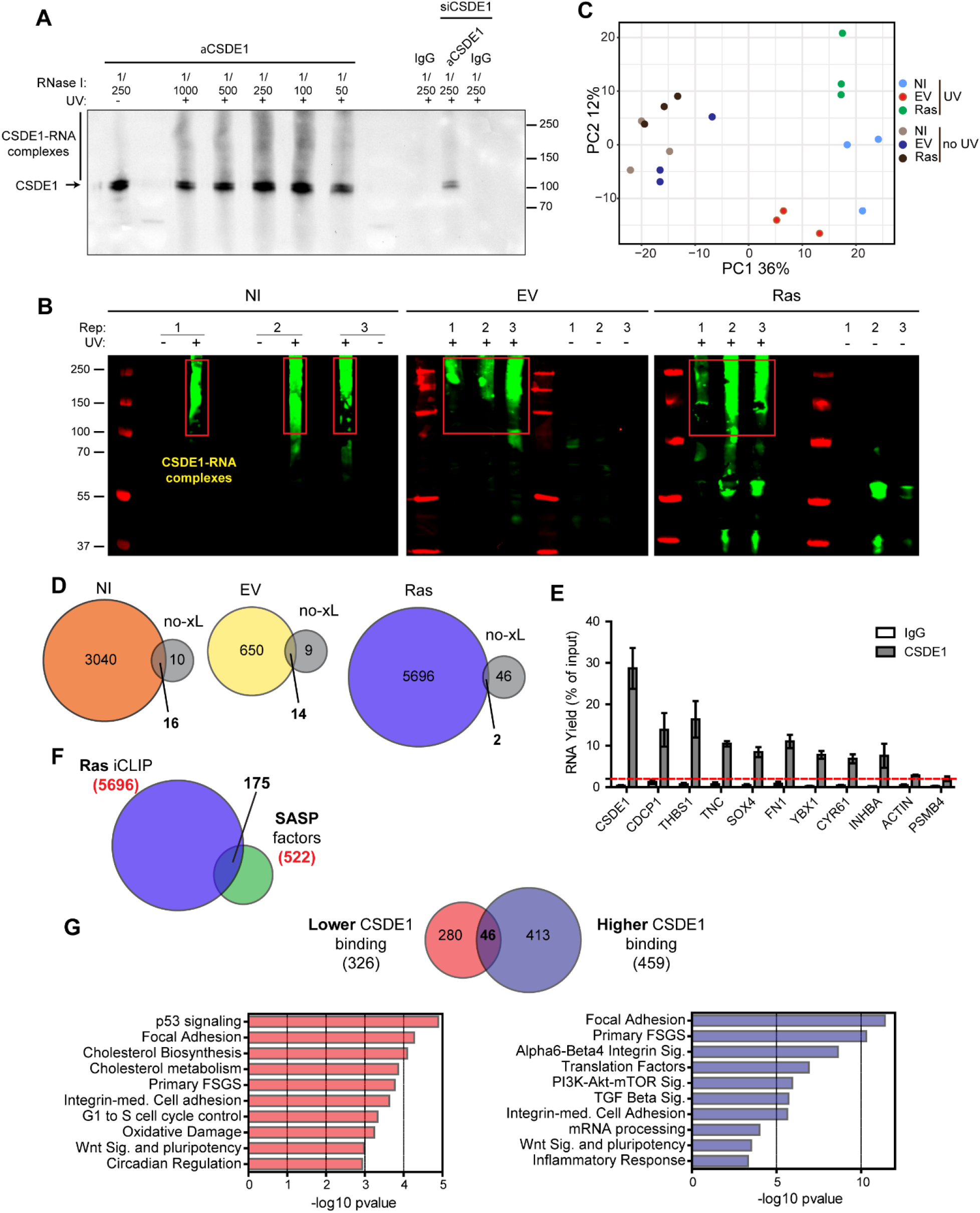
Identification of CSDE1 targets by iCLIP-seq (Related to Figure 3) **(A)** Specificity of CSDE1 immunoprecipitation and CSDE1-RNA complex formation, assessed by Western blot. A smear indicating CSDE1-RNA complexes is detected at different RNase I concentrations, while it is absent in the non-UV control. CSDE1 depletion as well as immunoprecipitation with non-specific IgG were carried as controls. **(B)** CSDE1-RNA complex formation in 3 biological replicates per condition using non-UV crosslinked samples as controls. Complexes were visualized by ligation of an infrared oligonucleotide used in the first step of library preparation, and visualized by fluorescence. Red squares highlight CSDE1-RNA complexes. **(C)** PCA analysis of iCLIP-seq samples. **(D)** Overlap between iCLIP targets detected in each condition and their respective negative controls. **(E)** Validation of CSDE1 targets by RT-qPCR in EV-infected PMK. Percentage of immunoprecipitated input is shown for both CSDE1 IP and control IgG. ACTIN and PSMB4 were used as negative controls. CSDE1 was used as positive control. **(F)** GO analysis of targets that bind differentially to CSDE1 in the EV vs Ras conditions.

**Figure S2.**
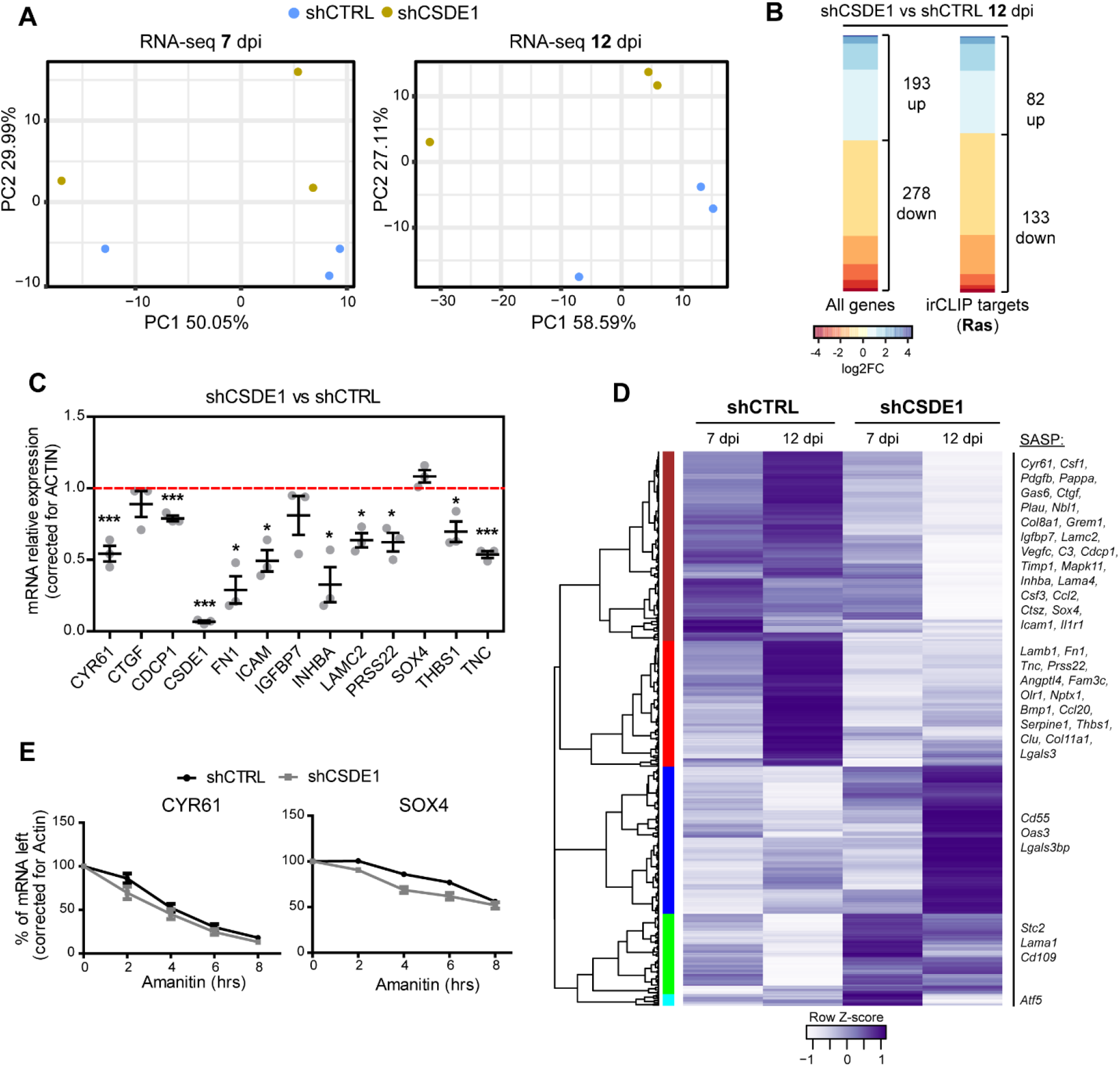
CSDE1 regulates SASP factor mRNA stability (Related to Figure 4) **(A)** PCA analysis of RNA-Seq samples at 7 and 12 dpi (n= 3 per condition). **(B)** Heatmaps of changes in the levels of all mRNAs or only CSDE1 targets at 12 dpi. **(C)** Validation of changes in SASP mRNA levels upon CSDE1 depletion (12 dpi) by RT-qPCR (n= 3). Significance was assessed by one sample t-test (*p-value <0.05, ***p-value < 0.001). Error bars represent SEM. **(D)** Unbiased hierarchical clustering of all changes after CSDE1 depletion. **(E)** The stability of CYR61 and SOX4 mRNAs does not significantly change upon CSDE1 depletion. Transcription in shCTRL and shCSDE1 cells was inhibited with alpha-amanitin, and mRNA levels tested at several time points thereafter by RT-qPCR (n= 5). The data were corrected for actin and normalized to the initial time point. Linear regression analysis was conducted to test the significance of differences between mRNA decay slopes. Error bars represent SEM.

**Figure S3.**
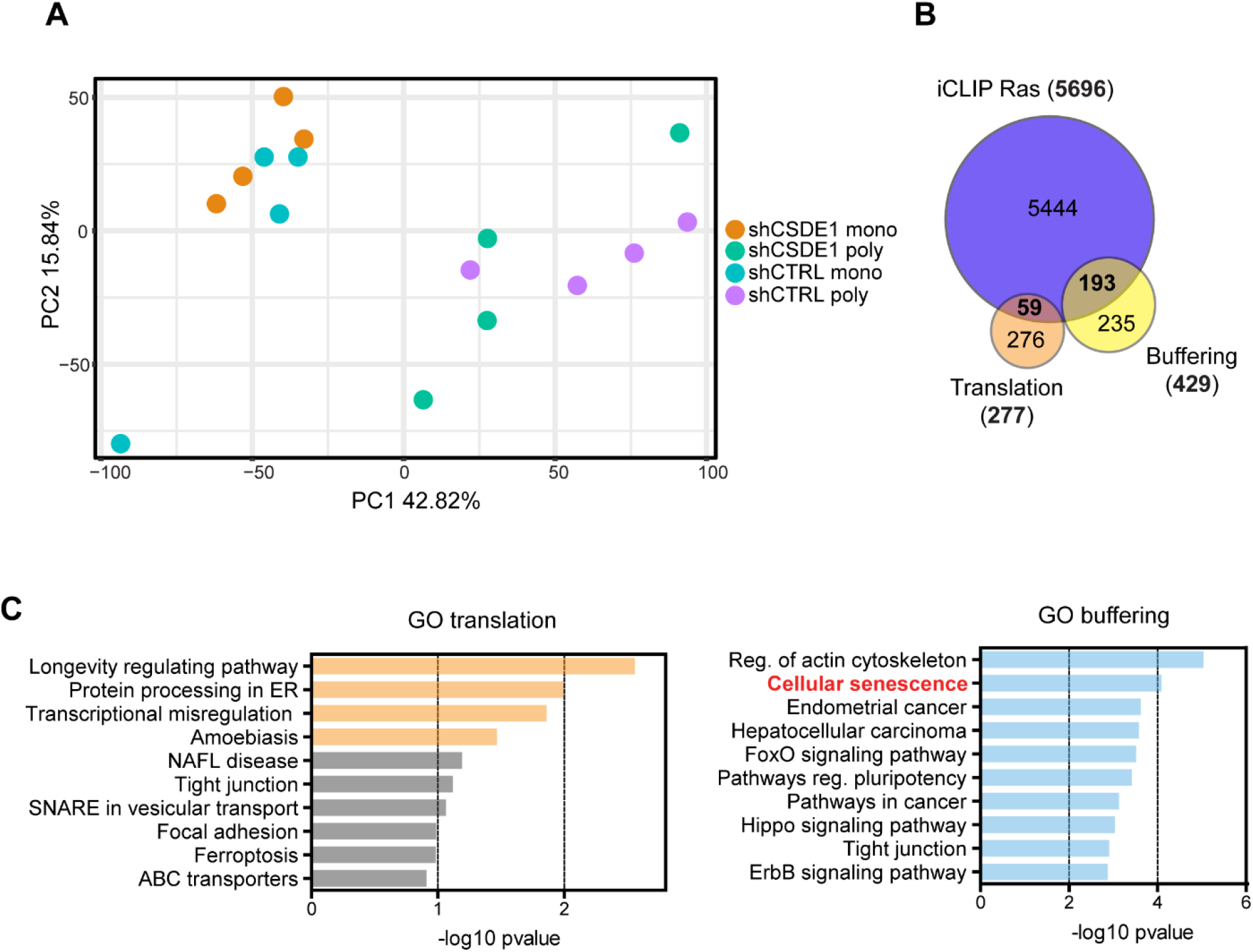
CSDE1 regulates translation of senescence-associated genes (Related to Figure 5) **(A)** PCA analysis of polysomal and monosomal RNA-seq samples. **(B)** Overlap between CSDE1 targets and transcripts that are translationally regulated or buffered upon CSDE1 depletion. **(C)** GO analysis of translationally regulated (left – orange indicates statistically significant enriched pathways) or buffered (right) transcripts within the CSDE1 target group.

**Figure S4.**
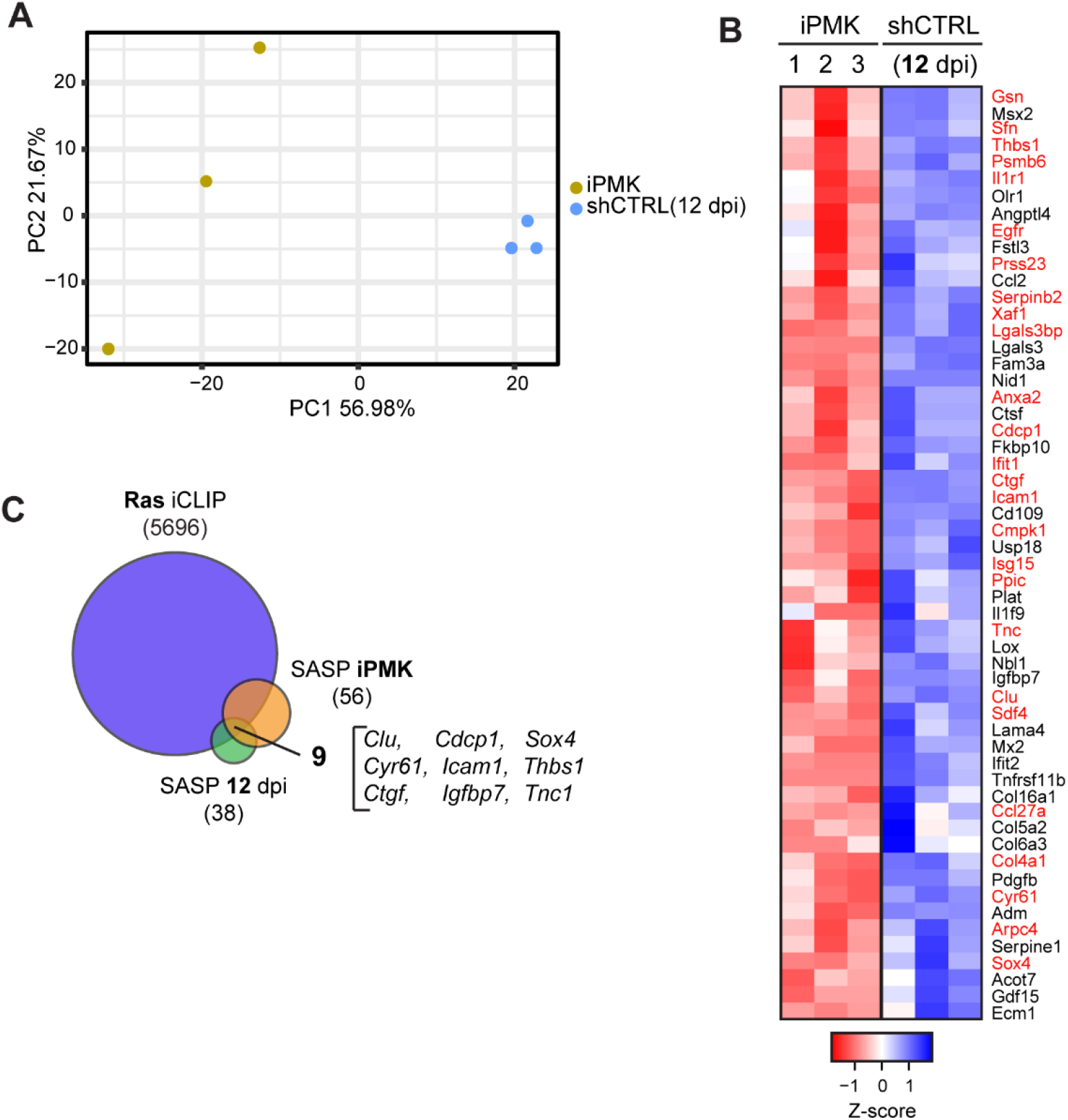
YBX1 depletion rescues the senescent phenotype (Related to Figure 6) **(A)** PCA analysis of iPMK and shCTRL (12 dpi) RNA-seq samples. **(B)** Heatmap of SASP components downregulated in iPMK cells compared to shCTRL (12 dpi). Half of them (red) are direct CSDE1 targets. **(C)** Overlap of CSDE1 targets in senescent cells (Ras) and SASP mRNAs downregulated upon CSDE1 depletion at 12 dpi and in iPMK.

## REFERENCES

Acosta, J.C., Banito, A., Wuestefeld, T., Georgilis, A., Janich, P., Morton, J.P., Athineos, D., Kang, T.W., Lasitschka, F., Andrulis, M., et al. (2013). A complex secretory program orchestrated by the inflammasome controls paracrine senescence. Nat Cell Biol 15, 978–990.

Anderson, E.C., and Catnaigh, P.O. (2015). Regulation of the expression and activity of Unr in mammalian cells. Biochem Soc Trans 43, 1241–1246.

Andrews, S. (2010). FastQC: A Quality Control Tool for High Throughput Sequence Data. Available online at: http://www.bioinformatics.babraham.ac.uk/projects/fastqc/.

Atlasi, Y., Jafarnejad, S.M., Gkogkas, C.G., Vermeulen, M., Sonenberg, N., and Stunnenberg, H.G. (2020). The translational landscape of ground state pluripotency. Nat Commun 11, 1617.

Bailey, T.L. (2011). DREME: motif discovery in transcription factor ChIP-seq data. Bioinformatics 27, 1653–1659.

Basisty, N., Kale, A., Jeon, O.H., Kuehnemann, C., Payne, T., Rao, C., Holtz, A., Shah, S., Sharma, V., Ferrucci, L., et al. (2020). A proteomic atlas of senescence-associated secretomes for aging biomarker development. PLoS Biol 18, e3000599.

Blevins, W.R., Tavella, T., Moro, S.G., Blasco-Moreno, B., Closa-Mosquera, A., Diez, J., Carey, L.B., and Alba, M.M. (2019). Extensive post-transcriptional buffering of gene expression in the response to severe oxidative stress in baker's yeast. Sci Rep 9, 11005.

Chang, N., Yi, J., Guo, G., Liu, X., Shang, Y., Tong, T., Cui, Q., Zhan, M., Gorospe, M., and Wang, W. (2010). HuR uses AUF1 as a cofactor to promote p16INK4 mRNA decay. Mol Cell Biol 30, 3875–3886.

Coppe, J.P., Rodier, F., Patil, C.K., Freund, A., Desprez, P.Y., and Campisi, J. (2011). Tumor suppressor and aging biomarker p16(INK4a) induces cellular senescence without the associated inflammatory secretory phenotype. J Biol Chem 286, 36396–36403.

Cunningham, F., Achuthan, P., Akanni, W., Allen, J., Amode, M.R., Armean, I.M., Bennett, R., Bhai, J., Billis, K., Boddu, S., et al. (2019). Ensembl 2019. Nucleic Acids Res 47, D745–D751.

Dobin, A., Davis, C.A., Schlesinger, F., Drenkow, J., Zaleski, C., Jha, S., Batut, P., Chaisson, M., and Gingeras, T.R. (2013). STAR: ultrafast universal RNA-seq aligner. Bioinformatics 29, 15–21.

Dormoy-Raclet, V., Markovits, J., Malato, Y., Huet, S., Lagarde, P., Montaudon, D., Jacquemin-Sablon, A., and Jacquemin-Sablon, H. (2007). Unr, a cytoplasmic RNA-binding protein with cold-shock domains, is involved in control of apoptosis in ES and HuH7 cells. Oncogene 26, 2595–2605.

El-Naggar, A.M., Veinotte, C.J., Cheng, H., Grunewald, T.G., Negri, G.L., Somasekharan, S.P., Corkery, D.P., Tirode, F., Mathers, J., Khan, D., et al. (2015). Translational Activation of HIF1alpha by YB-1 Promotes Sarcoma Metastasis. Cancer Cell 27, 682–697.

Elatmani, H., Dormoy-Raclet, V., Dubus, P., Dautry, F., Chazaud, C., and Jacquemin-Sablon, H. (2011). The RNA-binding protein Unr prevents mouse embryonic stem cells differentiation toward the primitive endoderm lineage. Stem Cells 29, 1504–1516.

Evdokimova, V., Tognon, C., Ng, T., Ruzanov, P., Melnyk, N., Fink, D., Sorokin, A., Ovchinnikov, L.P., Davicioni, E., Triche, T.J., et al. (2009). Translational activation of snail1 and other developmentally regulated transcription factors by YB-1 promotes an epithelial-mesenchymal transition. Cancer Cell 15, 402–415.

Fang, H., Yue, X., Li, X., and Taylor, J.S. (2005). Identification and characterization of high affinity antisense PNAs for the human unr (upstream of N-ras) mRNA which is uniquely overexpressed in MCF-7 breast cancer cells. Nucleic Acids Res 33, 6700–6711.

Fishbein, L., Leshchiner, I., Walter, V., Danilova, L., Robertson, A.G., Johnson, A.R., Lichtenberg, T.M., Murray, B.A., Ghayee, H.K., Else, T., et al. (2017). Comprehensive Molecular Characterization of Pheochromocytoma and Paraganglioma. Cancer Cell 31, 181–193.

Fregoso, O.I., Das, S., Akerman, M., and Krainer, A.R. (2013). Splicing-factor oncoprotein SRSF1 stabilizes p53 via RPL5 and induces cellular senescence. Mol Cell 50, 56–66.

Gorgoulis, V., Adams, P.D., Alimonti, A., Bennett, D.C., Bischof, O., Bishop, C., Campisi, J., Collado, M., Evangelou, K., Ferbeyre, G., et al. (2019). Cellular Senescence: Defining a Path Forward. Cell 179, 813–827.

Groisman, I., Ivshina, M., Marin, V., Kennedy, N.J., Davis, R.J., and Richter, J.D. (2006). Control of cellular senescence by CPEB. Genes Dev 20, 2701–2712.

Guo, A.X., Cui, J.J., Wang, L.Y., and Yin, J.Y. (2020). The role of CSDE1 in translational reprogramming and human diseases. Cell Commun Signal 18, 14.

Hanahan, D., and Weinberg, R.A. (2011). Hallmarks of cancer: the next generation. Cell 144, 646–674.

Hernandez-Segura, A., de Jong, T.V., Melov, S., Guryev, V., Campisi, J., and Demaria, M. (2017). Unmasking Transcriptional Heterogeneity in Senescent Cells. Curr Biol 27, 2652–2660 e2654.

Herranz, N., Gallage, S., Mellone, M., Wuestefeld, T., Klotz, S., Hanley, C.J., Raguz, S., Acosta, J.C., Innes, A.J., Banito, A., et al. (2015). mTOR regulates MAPKAPK2 translation to control the senescence-associated secretory phenotype. Nat Cell Biol 17, 1205–1217.

Herranz, N., and Gil, J. (2018). Mechanisms and functions of cellular senescence. J Clin Invest 128, 1238–1246.

Hollmann, N.M., Jagtap, P.K.A., Masiewicz, P., Guitart, T., Simon, B., Provaznik, J., Stein, F., Haberkant, P., Sweetapple, L.J., Villacorta, L., et al. (2020). Pseudo-RNA-Binding Domains Mediate RNA Structure Specificity in Upstream of N-Ras. Cell Rep 32, 107930.

Horos, R., Ijspeert, H., Pospisilova, D., Sendtner, R., Andrieu-Soler, C., Taskesen, E., Nieradka, A., Cmejla, R., Sendtner, M., Touw, I.P., et al. (2012). Ribosomal deficiencies in Diamond-Blackfan anemia impair translation of transcripts essential for differentiation of murine and human erythroblasts. Blood 119, 262–272.

Huber, W., Carey, V.J., Gentleman, R., Anders, S., Carlson, M., Carvalho, B.S., Bravo, H.C., Davis, S., Gatto, L., Girke, T., et al. (2015). Orchestrating high-throughput genomic analysis with Bioconductor. Nat Methods 12, 115–121.

Ju Lee, H., Bartsch, D., Xiao, C., Guerrero, S., Ahuja, G., Schindler, C., Moresco, J.J., Yates, J.R. 3rd, Gebauer, F., Bazzi, H., et al. (2017). A post-transcriptional program coordinated by CSDE1 prevents intrinsic neural differentiation of human embryonic stem cells. Nat Commun 8, 1456.

Kang, C., Xu, Q., Martin, T.D., Li, M.Z., Demaria, M., Aron, L., Lu, T., Yankner, B.A., Campisi, J., and Elledge, S.J. (2015). The DNA damage response induces inflammation and senescence by inhibiting autophagy of GATA4. Science 349, aaa5612.

Kotake, Y., Ozawa, Y., Harada, M., Kitagawa, K., Niida, H., Morita, Y., Tanaka, K., Suda, T., and Kitagawa, M. (2013). YB1 binds to and represses the p16 tumor suppressor gene. Genes Cells 18, 999–1006.

Kuleshov, M.V., Jones, M.R., Rouillard, A.D., Fernandez, N.F., Duan, Q., Wang, Z., Koplev, S., Jenkins, S.L., Jagodnik, K.M., Lachmann, A., et al. (2016). Enrichr: a comprehensive gene set enrichment analysis web server 2016 update. Nucleic Acids Res 44, W90–97.

Kwon, E., Todorova, K., Wang, J., Horos, R., Lee, K.K., Neel, V.A., Negri, G.L., Sorensen, P.H., Lee, S.W., Hentze, M.W., et al. (2018). The RNA-binding protein YBX1 regulates epidermal progenitors at a posttranscriptional level. Nat Commun 9, 1734.

Lee, J.H., Jung, M., Hong, J., Kim, M.K., and Chung, I.K. (2018). Loss of RNA-binding protein HuR facilitates cellular senescence through posttranscriptional regulation of TIN2 mRNA. Nucleic Acids Res 46, 4271–4285.

Li, H., Handsaker, B., Wysoker, A., Fennell, T., Ruan, J., Homer, N., Marth, G., Abecasis, G., Durbin, R., and Genome Project Data Processing, S. (2009). The Sequence Alignment/Map format and SAMtools. Bioinformatics 25, 2078–2079.

Liang, S., Bellato, H.M., Lorent, J., Lupinacci, F.C.S., Oertlin, C., van Hoef, V., Andrade, V.P., Roffe, M., Masvidal, L., Hajj, G.N.M., et al. (2018). Polysome-profiling in small tissue samples. Nucleic Acids Res 46, e3.

Liao, Y., Smyth, G.K., and Shi, W. (2019). The R package Rsubread is easier, faster, cheaper and better for alignment and quantification of RNA sequencing reads. Nucleic Acids Res 47, e47.

Liu, H., Li, X., Dun, M.D., Faulkner, S., Jiang, C.C., and Hondermarck, H. (2020). Cold Shock Domain Containing E1 (CSDE1) Protein is Overexpressed and can be Targeted to Inhibit Invasiveness in Pancreatic Cancer Cells. Proteomics, e1900331.

Love, M.I., Huber, W., and Anders, S. (2014). Moderated estimation of fold change and dispersion for RNA-seq data with DESeq2. Genome Biol 15, 550.

Majumder, M., House, R., Palanisamy, N., Qie, S., Day, T.A., Neskey, D., Diehl, J.A., and Palanisamy, V. (2016). Correction: RNA-Binding Protein FXR1 Regulates p21 and TERC RNA to Bypass p53-Mediated Cellular Senescence in OSCC. PLoS Genet 12, e1006411.

Martinez-Useros, J., Garcia-Carbonero, N., Li, W., Fernandez-Acenero, M.J., Cristobal, I., Rincon, R., Rodriguez-Remirez, M., Borrero-Palacios, A., and Garcia-Foncillas, J. (2019). UNR/CSDE1 Expression Is Critical to Maintain Invasive Phenotype of Colorectal Cancer through Regulation of c-MYC and Epithelial-to-Mesenchymal Transition. J Clin Med 8.

Maurelli, R., Tinaburri, L., Gangi, F., Bondanza, S., Severi, A.L., Scarponi, C., Albanesi, C., Mesiti, G., Guerra, L., Capogrossi, M.C., et al. (2016). The role of oncogenic Ras in human skin tumorigenesis depends on the clonogenic potential of the founding keratinocytes. J Cell Sci 129, 1003–1017.

Maurya, P.K., Mishra, A., Yadav, B.S., Singh, S., Kumar, P., Chaudhary, A., Srivastava, S., Murugesan, S.N., and Mani, A. (2017). Role of Y Box Protein-1 in cancer: As potential biomarker and novel therapeutic target. J Cancer 8, 1900–1907.

Mihailovich, M., Militti, C., Gabaldon, T., and Gebauer, F. (2010). Eukaryotic cold shock domain proteins: highly versatile regulators of gene expression. Bioessays 32, 109–118.

Mordovkina, D., Lyabin, D.N., Smolin, E.A., Sogorina, E.M., Ovchinnikov, L.P., and Eliseeva, I. (2020). Y-Box Binding Proteins in mRNP Assembly, Translation, and Stability Control. Biomolecules 10.

Nacarelli, T., Lau, L., Fukumoto, T., Zundell, J., Fatkhutdinov, N., Wu, S., Aird, K.M., Iwasaki, O., Kossenkov, A.V., Schultz, D., et al. (2019). NAD(+) metabolism governs the proinflammatory senescence-associated secretome. Nat Cell Biol 21, 397–407.

Oertlin, C., Lorent, J., Murie, C., Furic, L., Topisirovic, I., and Larsson, O. (2019). Generally applicable transcriptome-wide analysis of translation using anota2seq. Nucleic Acids Res 47, e70.

Patel, G.P., Ma, S., and Bag, J. (2005). The autoregulatory translational control element of poly(A)-binding protein mRNA forms a heteromeric ribonucleoprotein complex. Nucleic Acids Res 33, 7074–7089.

R Core, T. (2019). R: A Language and Environment for Statistical Computing.

Radine, C., Peters, D., Reese, A., Neuwahl, J., Budach, W., Janicke, R.U., and Sohn, D. (2020). The RNA-binding protein RBM47 is a novel regulator of cell fate decisions by transcriptionally controlling the p53-p21-axis. Cell Death Differ 27, 1274–1285.

Ramirez, F., Ryan, D.P., Gruning, B., Bhardwaj, V., Kilpert, F., Richter, A.S., Heyne, S., Dundar, F., and Manke, T. (2016). deepTools2: a next generation web server for deep-sequencing data analysis. Nucleic Acids Res 44, W160–165.

Rao, S.G., and Jackson, J.G. (2016). SASP: Tumor Suppressor or Promoter? Yes! Trends Cancer 2, 676–687.

Ritchie, M.E., Phipson, B., Wu, D., Hu, Y., Law, C.W., Shi, W., and Smyth, G.K. (2015). limma powers differential expression analyses for RNA-sequencing and microarray studies. Nucleic Acids Res 43, e47.

Ritschka, B., Storer, M., Mas, A., Heinzmann, F., Ortells, M.C., Morton, J.P., Sansom, O.J., Zender, L., and Keyes, W.M. (2017). The senescence-associated secretory phenotype induces cellular plasticity and tissue regeneration. Genes Dev 31, 172–183.

Robinson, M.D., and Oshlack, A. (2010). A scaling normalization method for differential expression analysis of RNA-seq data. Genome Biol 11, R25.

Ruiz, L., Traskine, M., Ferrer, I., Castro, E., Leal, J.F., Kaufman, M., and Carnero, A. (2008). Characterization of the p53 response to oncogene-induced senescence. PLoS One 3, e3230.

Ruscetti, M., Morris, J.P.t., Mezzadra, R., Russell, J., Leibold, J., Romesser, P.B., Simon, J., Kulick, A., Ho, Y.J., Fennell, M., et al. (2020). Senescence-Induced Vascular Remodeling Creates Therapeutic Vulnerabilities in Pancreas Cancer. Cell 181, 424–441 e421.

Sanduja, S., Kaza, V., and Dixon, D.A. (2009). The mRNA decay factor tristetraprolin (TTP) induces senescence in human papillomavirus-transformed cervical cancer cells by targeting E6-AP ubiquitin ligase. Aging (Albany NY) 1, 803–817.

Schepens, B., Tinton, S.A., Bruynooghe, Y., Parthoens, E., Haegman, M., Beyaert, R., and Cornelis, S. (2007). A role for hnRNP C1/C2 and Unr in internal initiation of translation during mitosis. EMBO J 26, 158–169.

Serrano, M., Lin, A.W., McCurrach, M.E., Beach, D., and Lowe, S.W. (1997). Oncogenic ras provokes premature cell senescence associated with accumulation of p53 and p16INK4a. Cell 88, 593–602.

Tian, N.Y., Qi, Y.J., Hu, Y., Yin, B., Yuan, J.G., Qiang, B.Q., Peng, X.Z., and Han, W. (2018). RNA-binding Protein UNR Promotes Glioma Cell Migration and Regulates the Expression of Ribosomal Protein L9. Chin Med Sci J 33, 143–151.

Wang, B., Kohli, J., and Demaria, M. (2020). Senescent Cells in Cancer Therapy: Friends or Foes? Trends Cancer 6, 838–857.

Wang, W. (2012). Regulatory RNA-binding proteins in senescence. Ageing Res Rev 11, 485–490.

Wang, W., Martindale, J.L., Yang, X., Chrest, F.J., and Gorospe, M. (2005). Increased stability of the p16 mRNA with replicative senescence. EMBO Rep 6, 158–164.

Wang, W., Yang, X., Cristofalo, V.J., Holbrook, N.J., and Gorospe, M. (2001). Loss of HuR is linked to reduced expression of proliferative genes during replicative senescence. Mol Cell Biol 21, 5889–5898.

Warnes, G.R. (2015). gplots: various R programming tools for plotting data.

Wickham, H. (2016). ggplot2: Elegant Graphics for Data Analysis. (Springer-Verlag New York).

Wurth, L., Papasaikas, P., Olmeda, D., Bley, N., Calvo, G.T., Guerrero, S., Cerezo-Wallis, D., Martinez-Useros, J., Garcia-Fernandez, M., Huttelmaier, S., et al. (2016). UNR/CSDE1 Drives a Post-transcriptional Program to Promote Melanoma Invasion and Metastasis. Cancer Cell 30, 694–707.

Zhu, H., Blake, S., Kusuma, F.K., Pearson, R.B., Kang, J., and Chan, K.T. (2020). Oncogene-induced senescence: From biology to therapy. Mech Ageing Dev 187, 111229.

Jensen, K.B., Driskell, R.R., and Watt, F.M. (2010). Assaying proliferation and differentiation capacity of stem cells using disaggregated adult mouse epidermis. Nat Protoc 5, 898–911.

